# Loss of tissue non-specific alkaline phosphatase (TNAP) enzyme activity in cerebral microvessels is coupled to persistent neuroinflammation and behavioral deficits in late sepsis

**DOI:** 10.1101/732008

**Authors:** Divine C. Nwafor, Sreeparna Chakraborty, Allison L. Brichacek, Sujung Jun, Catheryne A. Gambill, Wei Wang, Elizabeth B. Engler-Chiurazzi, Duaa Dakhlallah, Stanley A. Benkovic, Candice M. Brown

## Abstract

Sepsis is characterized as a host response to systemic inflammation and infection that may lead to multi-organ dysfunction and eventual death. While acute brain dysfunction is common among all sepsis patients, chronic neurological impairment is prevalent among sepsis survivors. The brain microvasculature has recently emerged as a major determinant of sepsis-associated brain dysfunction, yet the mechanisms that underlie its associated neuroimmune perturbations and behavioral deficits are not well understood. A growing body of data suggests that the loss of tissue nonspecific alkaline phosphatase (TNAP) enzyme activity in cerebral microvessels may be associated with changes in endothelial cell barrier integrity. The objective of this study was to determine the important mechanisms linking alterations in cerebrovascular TNAP enzyme activity to underlying neurological impairment in late sepsis. We hypothesized that the disruption of TNAP enzymatic activity in cerebral microvessels would be coupled to the sustained loss of brain microvascular integrity, elevated neuroinflammatory responses, and behavioral deficits. Male mice were subjected to cecal ligation and puncture (CLP), a model of experimental sepsis, and assessed up to seven days post-sepsis. All mice were observed daily for sickness behavior and underwent behavioral testing. Our results showed a significant decrease in brain microvascular TNAP enzyme activity in the somatosensory cortex and spinal cord of septic mice but not in the CA1 and CA3 hippocampal regions. Analyses of whole brain myeloid and T-lymphoid cell populations also revealed a persistent elevation of infiltrating leukocytes, which included both neutrophil and monocyte myeloid derived suppressor cells (MDSCs). Regional analyses of the somatosensory cortex, hippocampus, and spinal cord revealed significant astrogliosis and microgliosis in the cortex and spinal cord of septic mice that was accompanied by significant microgliosis in the CA1 and CA3 hippocampal regions. Assessment of behavioral deficits revealed no changes in learning and memory or evoked locomotion. However, the hot plate test uncovered a novel anti-nociceptive phenotype in our septic mice, and we speculate that this phenotype may be a consequence of sustained GFAP astrogliosis and loss of TNAP activity in the somatosensory cortex of septic mice. Taken together, these results demonstrate that the loss of TNAP enzyme activity in cerebral microvessels during late sepsis is coupled to sustained neuroimmune dysfunction which may underlie, in part, the chronic neurological impairments observed in sepsis survivors.

**Highlights:**

- Alkaline phosphatase activity in brain microvessels is diminished in sepsis.
- Loss of alkaline phosphatase activity is coupled to the loss of barrier integrity.
- Brain infiltration of myeloid and T-lymphoid cells persists in late sepsis.
- Septic mice exhibit a novel anti-nociceptive phenotype.
- Cortical astrogliosis and microgliosis persist in late sepsis.

## 1. Introduction

Sepsis is a potentially fatal clinical syndrome that results from an excessive systemic inflammatory response to an infection (Stare et al., 2015; Zaghloul et al., 2017). This syndrome accounts for 30 million cases annually, and about 6 million deaths worldwide (Fleischmann et al., 2016). A unique feature of sepsis is that it is often comorbid or may precipitate from other disease conditions such as stroke, cancer, or diabetes. (Boehme, Ranawat, Luna, Kamel, & Elkind, 2017; Nwafor et al., 2019; H. E. Wang et al., 2012). It is anticipated that the rise in these chronic diseases will increase the global prevalence of sepsis (Ovbiagele et al., 2013; Stoller et al., 2016). Yet, therapeutic strategies to treat septic patients remain limited, as there are no FDA-approved drugs to treat sepsis, and the primary treatments are largely limited to anti-microbial drugs, supportive fluids, and vasopressors (Fink & Warren, 2014). In addition, the mechanisms underlying the long-term consequences of sepsis on the brain and periphery remain elusive (Iskander et al., 2013; Iwashyna, Ely, Smith, & Langa, 2010).

Sepsis survivors are burdened with an increased risk of infections, sensorimotor abnormalities, and cognitive decline (Iwashyna et al., 2010; Nwafor et al., 2019; Zauner et al., 2002). The central nervous system (CNS), particularly the brain, is severely affected in sepsis and sepsis-associated cognitive impairment is an independent predictor of mortality (Gofton & Young, 2012). Persistent peripheral inflammation in sepsis contributes to a dysfunctional brain microvasculature and facilitates the trafficking of peripheral immune cell populations, cytokines and neurotoxins into the brain which, collectively, promote neuroinflammation and alter brain function (Andonegui et al., 2018; Singer et al., 2016). Short-term brain microvascular dysfunction in murine models of experimental sepsis has been demonstrated by increased barrier permeability and diminished tight junction (TJ) proteins 24 h post-sepsis (Towner et al., 2018; P. Wang, Hu, Yao, & Li, 2018). Thus, a better understanding of the cellular and molecular mechanisms that contribute to brain microvascular dysfunction in sepsis may provide insights to mitigate and improve long term neurological outcomes in sepsis survivors.

We and others have shown that the ectoenzyme known as tissue non-specific alkaline phosphatase (TNAP) may play a critical role in brain microvascular function and dysfunction (Deracinois, Lenfant, Dehouck, & Flahaut, 2015; Nwafor et al., 2019). TNAP, also known as bone/liver/kidney alkaline phosphatase, is one of four alkaline phosphatase genes in the mammalian genome (Betz, Firth, & Goldstein, 1980; Street et al., 2013; Vorbrodt, Lossinsky, & Wisniewski, 1986). Peripheral TNAP enzymatic activity, most likely generated within neutrophils, liver, and kidneys, has been shown to play an important role in attenuating inflammation and improving survival in sepsis rodent models (Bender et al., 2015; Verweij et al., 2004). TNAP enzyme activity is also highly elevated in cerebral microvessels and has been used as a histological marker of the brain microvasculature for nearly a century (Betz et al., 1980; Friede, 1966; Vorbrodt et al., 1986). While a clearly delineated physiological role for TNAP at the brain microvascular interface remains less understood, recent studies suggest a role for TNAP in the maintenance of brain endothelial cell integrity. An *in vitro* study demonstrated that inhibition of TNAP activity on brain endothelial cells by levamisole, a pan-AP enzyme inhibitor, worsened barrier function and increased cellular permeability (Deracinois et al., 2015). In a recent *in vivo* study, we reported that brain TNAP enzyme activity is decreased 24 h post-sepsis compared to sham-injured mice (Nwafor et al., 2019). Taken together, both findings demonstrate a plausible role for TNAP in maintaining brain microvascular barrier integrity during inflammation. However, it remains unclear whether the observed decrease in TNAP’s enzyme activity persists beyond 24 hours and whether specific brain regions exhibit selective changes in TNAP activity compared to others. The delineation between early (≤ 24 h) and late sepsis (> 36 h) is particularly important considering that most sepsis-related deaths occur in the late/hypo-inflammatory phase of sepsis (Otto et al., 2011).

The objective of this study was to determine the neuroimmune and behavioral changes which paralleled the loss of TNAP in cerebral microvessels in late sepsis, i.e. up to seven days post-sepsis. We incorporated an established a 7-day model of late sepsis, cecal ligation and puncture (CLP) (Crowell, Phillips, Kelleher, Soybel, & Lang, 2017; Vachharajani et al., 2014; X. Wang et al., 2016). We assessed novel behavioral and neuroinflammatory outcomes in late sepsis and coupled these outcomes to distinct alterations in the brain microvasculature. Our results revealed a loss of TNAP enzymatic activity in cerebral microvessels and diminished tight junctions, indicating a loss of barrier integrity. Enhanced cortical astrogliosis and microgliosis was accompanied by sustained leukocyte infiltration, and we also identified a novel cell population with characteristics of myeloid-derived suppressor cells. Assessment of behavioral deficits in late sepsis uncovered a novel anti-nociceptive phenotype in the absence of an impairment in learning and memory. Taken together, these results demonstrate that late sepsis is characterized by a novel neuroimmune phenotype that embodies loss of sustained loss of cerebral microvascular TNAP activity and persistent neuroinflammation.

## 2. Materials and Methods

### 2.1. Animals

All experiments were conducted in accordance with the National Institutes of Health Guide for the Care and Use of Laboratory Animals and were approved by the Institutional Animal Care and Use Committee at West Virginia University. Male wild-type (WT; *C57BL6/J*) mice were bred in West Virginia University Health Sciences Center vivarium facilities and used for experiments at 3-5 months old. All mice were generated from C57BL/6J breeding pairs obtained from Jackson Labs (Bar Harbor, ME, Catalog # 000664). Mice were group housed in environmentally controlled conditions with a reverse light cycle (12:12 h light/dark cycle at 21 ± 1°C) and provided food and water *ad libitum*.

### 2.2. Cecal ligation and puncture (CLP)

The cecal ligation and puncture (CLP) model of moderate polymicrobial sepsis was employed as previously described (Vachharajani et al., 2014). Briefly, mice were anesthetized by the inhalation of 1-2% isoflurane. Abdominal access was obtained via a midline incision following topical application of 2% lidocaine (Hospira, Lake Forest, IL). The cecum was isolated and ligated with a 4-0 silk ligature below the ileocecal valve and then punctured twice with a 22 G needle through and through. Fecal matter (approximately 1 mm) was extruded from the hole, and the cecum was placed back into the abdominal cavity. The abdominal muscle layer was closed with 6-0 sutures (Ethilon, Cornelia, Georgia), while the abdominal skin layer was closed with 5-0 sutures (Ethilon, Cornelia, Georgia). Sham-operated animals had their cecum isolated and then returned to the peritoneal cavity without being ligated or punctured. One mL of sterile 0.9% saline was administered subcutaneously after sepsis procedure for fluid resuscitation in sham- and CLP-operated mice. All mice were housed according to treatment group and placed on heating pads for 3 days following sepsis procedure. Mice were monitored for survival and sepsis-associated clinical signs twice daily for 6 days and underwent a series of behavioral assessments.

### 2.3. Modified murine sepsis score (MMSS)

We modified a validated murine sepsis severity score (MSS) (Shrum et al., 2014) to examine eight sickness behavior components: appearance, weight, activity, response to stimulus, eyes, posture, diarrhea, and respiratory rate, henceforth referred to as the modified murine sepsis severity score (MMSS). Each component is scored from 0 (normal/no change) to 4 (severe change), and the MMSS is the grand total of all eight components collected each day post-sepsis over a 6-day period for each animal. The average percent weight loss score was derived from the MMSS and defined on a scale from 0 to 4 (i.e. a score of 0=0-5%, 1=5.1-10%, 2=10.1-15%, 3=15.1-20%, and 4=>20.1% average percent weight loss). Animals with a MMSS greater than 24 were euthanized and excluded from behavioral assessments.

### 2.4. Tissue collecting and processing

Mice were deeply anesthetized with isoflurane and perfused intracardially with a perfusion pump (Masterflex 7524-10, Cole-Parmer, Vernon Hills, IL) set to 5.0 mL/min. Blood was removed with 25 mL of 0.9% saline followed by perfusion and fixation with 50 mL 4% chilled paraformaldehyde (PFA, Fisher Scientific, Pittsburgh, PA). Perfused brains and spinal cords were removed from the skull and spinal canal, respectively, and post-fixed in 4% PFA overnight at 4°C. On the following day, brains and spinal cords were rinsed in 0.01 M phosphate buffered saline (PBS) and incubated sequentially in 10%, 20%, and 30% sucrose in PBS for 24 h each. Following successive incubations in sucrose, brains and spinal cords were co-embedded in a 15% gelatin matrix in groups of twelve per matrix for simultaneous sectioning. The gelatin block was processed sequentially through 4% PFA for 24 h, 15% sucrose for 24 h, and 30% sucrose for 48 h. The block was trimmed and placed in a −80^°^C freezer for 30 min. Sectioning was performed in the coronal plane at 30 µm on a sliding microtome (HM 450, ThermoFisher Scientific, Waltham, MA) equipped with a 3×3 freezing stage (BFS-40MPA, Physitemp, Clifton, NJ) at −20°C. Sections were collected into a series of six cups filled with PBS/0.06% sodium azide. Adjacent cups were used for sequential histological staining or immunostaining.

### 2.5. Immunohistochemistry

Following sequential cuts of the gelatin blocks on the microtome, sections were immunostained using standard free-floating immunohistochemistry techniques as described (Bachman, 2013). Briefly, all sections were blocked with 80 mL PBS, 10 mL of methanol (Fisher Scientific, Pittsburgh, PA), and 10 mL of 30% hydrogen peroxide (Fisher Scientific, Pittsburgh, PA) and incubated on a shaker (Model 55D, Reliable Scientific, Hernando, MS) for 15 min. Sections were then washed 3 times and permeabilized for 30 min on a shaker with 1.83% lysine (Fisher Scientific, Pittsburgh, PA) in 1% Triton (Sigma-Aldrich, St. Louis, MO), and 4% heat-inactivated horse serum (Sigma-Aldrich, St. Louis, MO). Sections were then incubated for 24 h with primary antibodies at room temperature, followed by a 2 h incubation with the appropriate secondary antibody at room temperature. The following primary antibodies were used with working dilutions indicated in parentheses: Iba-1 (Invitrogen (1:1000), Carlsbad, CA), GFAP (Agilent (1:10,000), Santa Clara, CA), Claudin 5 (GeneTex (1:500), Irvine, CA), TMEM 119 (Abcam (1:1000), Cambridge, MA), NeuN (Cell Signaling Technologies (1:500), Danvers, MA), and choline acetyltransferase (ChAT) antibody (Abcam (1:1000), Cambridge, MA).

### 2.6. Tissue non-specific alkaline phosphatase (TNAP) enzyme histology

Brains and spinal cords were evaluated for alkaline phosphatase enzyme activity by staining free-floating with a BCIP/NBT substrate kit (SK-5400, Vector Laboratories, Burlingame, CA) as previously described (Nwafor et al., 2019). Sections were rinsed in Dulbecco’s PBS (DPBS) twice for 5 min each, once in 0.1M Tris-HCl (pH = 9.5) for 5 min and incubated in the staining solution for 8 h at room temperature. Following three rinses in 0.01 M PBS, the sections were mounted onto microscope slides (Unifrost+, Azer Scientific, Morgantown, PA), air-dried overnight, dehydrated through a standard dehydration series, and cover-slipped with Permount (Fisher Scientific, Pittsburgh, PA).

### 2.7. Image analysis

Sections were viewed on a Leica DM6B microscope (Leica Camera, Allendale, NJ) and images were captured using Leica LASX software (Leica Microsystems, Buffalo Grove, IL). All brain regions of interest i.e. the striatum, CA1, CA3, somatosensory cortex, and basal forebrain (diagonal band of Broca and medial septum) were identified by referring to the Allen Institute Brain Atlas (http://mouse.brain-map.org). Analysis of Iba-1 and TMEM 119 was conducted in the somatosensory cortex (20X (TMEM119), and 20X and 40X (Iba-1) magnifications), CA1 (40X (TMEM119 and Iba-1) magnification), and CA3 brain regions (40X (TMEM119 and Iba-1) magnification). Analysis for GFAP was conducted in the somatosensory cortex (20X magnification), CA1 (40X magnification), and CA3 (40X magnification). Cell counting analysis for NeuN was conducted in the CA1 and CA3 hippocampal regions at 40X magnification. Cholinergic (ChAT-positive) neurons were analyzed in the basal forebrain region at 40X magnification. Analysis for TNAP enzyme activity was conducted in the somatosensory cortex at 20X magnification. Three random images were collected per animal per brain region of interest. Spinal cords were analyzed for GFAP, Iba-1, and TNAP. All collected images were converted to an 8-bit image and quantified in FIJI/Image J version 2.0 software.

### 2.8. Flow cytometric sample preparation, gating strategy and analysis

Single cell suspensions were obtained as previously described using the Miltenyi adult brain dissociation kit (Miltenyi Biotec, Auburn, CA) according to the manufacturer’s instructions (Garcia-Bonilla et al., 2018; Vachharajani et al., 2014). Briefly, mice were deeply anesthetized with isoflurane and were perfused intracardially with a perfusion pump (Masterflex 7524-10, Cole-Parmer, Vernon Hills, IL) set to 5.0 ml/min. Blood was removed with 25 mL 0.9% saline. Perfused whole brains were removed from the skull, minced, and homogenized mechanically into a Gentle MACS C tube containing 50 µL enzyme P + 1900 µL buffer Z and 20 µL of Buffer Y + 10 µL of Enzyme A (Miltenyi Biotec, Auburn, CA) and passed through a 70 µm cell strainer (Millipore, Temecula, CA) to obtain a single cell suspension. Cells were then collected by centrifugation at 300 x *g* for 10 min at 4°C. Red blood cells were lysed with 4 ml of ACK Lysis Buffer (Lonza, Walkersville, MD) per brain, incubated for 2-3 min at room temperature and washed. Viability and total brain cell yield was determined by trypan blue exclusion and cells were re-suspended to a final concentration of 2.5-3 × 10^6^ cells/ml with FACS buffer (0.01 M PBS, 5mM EDTA, 2% FBS). Cells were washed twice in cold 0.01M PBS and stained with fixable viability dye eFluor780 (eBioscience, San Diego, USA) for 30 min at 4°C in the dark. Cells were then briefly washed with FACS buffer and blocked with Ultra-Leaf purified anti-mouse CD16/32 (BioLegend, San Diego, CA) for 20 min. Following non-specific blocking, cells were stained with monoclonal antibodies for CD45-PE, CD11b-VioBlu, CD11c-PerCp-Vio700, Ly6G-APC, Ly6C-PE-Vio770 (Miltenyi Biotec, Auburn, CA) or CD4-FITC, CD25-PE, CD19-PerCp-Cy5.5 and CD8a-eFluor450 (eBioscience, San Diego, USA) for 10 min at 4°C. Appropriate single stained controls were prepared for fluorophore compensation using compensation beads (Invitrogen, Carlsbad, CA). For intracellular FOXP3 detection cells were fixed/permeabilized using a mouse regulatory T cell staining kit (eBioscience) after staining the surface markers (CD4 or CD25, Invitrogen, Carlsbad, CA) and stained with anti-FOXP3-PE-Cy5. Fluorescence was measured using a BD LSR Fortessa with FACS Diva software (BD Biosciences, San Jose, CA). All data were compensated and spectral overlap was minimized using automatic compensation method of BD FACS diva software (BD Biosciences, San Jose, CA).

Further analysis was carried in FCS Express 6.0 (De Novo Software, Glendale, CA). Single cells were identified by forward scatter and side scatter, and viable cells were gated for further analysis of positive cell populations. Viable cells were gated for CD45^+^ populations and then divided into T-lymphoid cells, which included cytotoxic T-cells (CD45^hi^, CD3^+^, CD8^+^), helper T-cells (CD45^hi^, CD3^+^,CD4^+^), and T-regulatory cells (CD4^+^,CD25^+^, Foxp3^+^); myeloid cells, which included neutrophils (CD45^hi^, CD11b^+^, Ly6C^-^, Ly6G^+^), monocytes (CD45^hi^, CD11c^+^, Ly6C^+^, Ly6G^-^), polymorphonuclear myeloid-derived suppressor cells (PMN-MDSCs) (CD45^hi^, CD11b^+^, Ly6C^low^, Ly6G^+^), and monocytic myeloid-derived suppressor cells (M-MDSCs) (CD45^hi^, CD11b^+^, Ly6C^hi^, Ly6G^-^); and brain residential microglia (CD45^low^, CD11b^+^, CD11c^-^). The quantified results represent the percentage of positive cells from the total CD45^+^ live cells.

### 2.9. Behavioral testing

#### 2.9.1. Open field (OF)

The open field testing paradigm was used to evaluate passive locomotor activity as previously described (Doll et al., 2015) and was administered on day 2 post-sepsis. A white plastic box (60 cm × 60 cm × 15 cm) was placed upon a table accompanied by five 60-watt lamps that provided indirect illumination. At the start of the trial, each mouse was placed at the center of the box and received a 1 h session recorded over 5 min intervals. Locomotor activity was recorded using a 4×8 photobeam activity system by San Diego Instruments (San Diego, CA). The dependent variables are the total horizontal movements (x, y direction) and rearing movements (z direction) over a 60 min period.

#### 2.9.2. Accelerating rotarod (AR)

The accelerating rotarod test was used to evaluate evoked locomotor performance and coordination as previously described (Doll et al., 2015) and was administered on day 3 post-sepsis. A textured plastic horizontal rod (3 cm in diameter) was mounted 14.5 cm above a pressure-sensitive base (Ugo Basile, Gemonio, Italy). For each trial, the mouse was placed on a moving rod that accelerates from 4 rpm to 44 rpm when the operator hits the start button. Acceleration is continued until the mouse fell onto the padded base or until 300 s elapsed. Latency to fall (s) onto the padded base was the dependent variable. All mice received 4 trials on the rotarod.

#### 2.9.3. Passive avoidance (PA)

The passive avoidance test is used to assess non-spatial aversive learning and memory (Barichello et al., 2005) and was administered days 4 and 5 post-sepsis. The test is performed using a two-compartment (illuminated and darkened) chamber connected by an automated door (Med Associates, VT). All animals undergo 3 trials (120 s acquisition, 15 min immediate retention, and 24 h retention). During the 120 s acquisition, each animal is placed in the illuminated compartment and receives a 0.3mA shock when it enters the darkened compartment. For the 300 s retention trials, the animal is placed in the illuminated chamber; however, no shock is initiated upon entry into darkened compartment. The dependent variable is the latency to enter the darkened compartment (s).

#### 2.9.4. Hot plate (HP)

The hot plate test is used to assess nociception (Vachon et al., 2013) and was administered on day 5 post-sepsis. Mice were individually placed on a preheated 55°C hot plate (IITC Life Science, CA), inside an open-ended cylindrical Plexiglas tube with a diameter of 30 cm. The latency (s) to respond to thermal stimuli and the total number of nociceptive behaviors (jumps as well as hind limb-lick, flick or flexion) exhibited over a 30 s time frame were measured. To prevent tissue damage, mice were removed from the hot plate after 30 s regardless of their response.

#### 2.9.5. Two-day radial arm water maze (2D-RAWM)

The 2D-RAWM was used as previously described (Colton et al., 2014; Kan et al., 2015) with minor modifications. Briefly, a 6-arm maze is submerged in a pool of opaque water, and an invisible platform placed at the end of 1 arm below the surface of the water. Each mouse received 15 trials per day for 2 days and the start arm was changed on each trial, however, the goal arm remained the same for all animals. Using static visual cues, the mouse learned the position of the platform. The number of errors (incorrect arm entries) is counted across 1 min trials. If a mouse failed to locate the platform in 1 min, the mouse was guided to the platform by the investigator and allowed to stay on the platform for 15 s before returning the mouse to a heated cage. The number of errors is averaged over 3 trials per block, resulting in 5 blocks (15 trials) per day per animal, and a total of 10 blocks (30 trials) for the two-day period. The latency to get to the platform (s) and the number of errors are the dependent variables measured. The 2D-RAWM was performed on days 5 (acquisition) and 6 (retention) post-sepsis. It is important to note that mice that participated in the hot plate test and the passive avoidance test did not participate in the 2D-RAWM and vice versa due to the stressful impact of both tests on the mice. To avoid any potential influence of aversive testing experience, a separate cohort of mice was used for PA/HP testing than those that underwent 2D-RAWM.

### 2.10. Experimental design and statistical analysis

All experiments were executed to enhance rigor and avoid experimenter bias according to ARRIVE guidelines (Kilkenny et al., 2010). Only male mice were used for all experiments because males are more susceptible to sepsis than females as previously published (Angele, Pratschke, Hubbard, & Chaudry, 2014). The use of male mice also allows for comparisons with a number of published studies which address similar outcomes (Andonegui et al., 2018; Denstaedt et al., 2018; Zaghloul et al., 2017). Behavioral assays were conducted from the least aversive (i.e. open field) to most aversive (i.e. 2D-RAWM). The investigator was blinded to the treatment groups for all behavior tests and image analysis. Animals that died 24 h after a behavior test was conducted were excluded from the study. See Fig. 1 for an experimental timeline and outline of behavioral assays performed. All statistical analyses were conducted in GraphPad Prism 8.1 (GraphPad Software, La Jolla, CA). Results are expressed as mean ± SEM and *p*-values ≤0.05 were considered significant. Survival was analyzed using a log-rank test. All other results were analyzed using a two-tailed unpaired Student’s t-test or a repeated two-way analysis of variance (ANOVA). Datasets that did not display a Gaussian distribution were subjected to a comparable non-parametric analysis as indicated in the text and figure legends. All *p*-values and *n* values are indicated in the text and figure legends.

**Figure 1.**
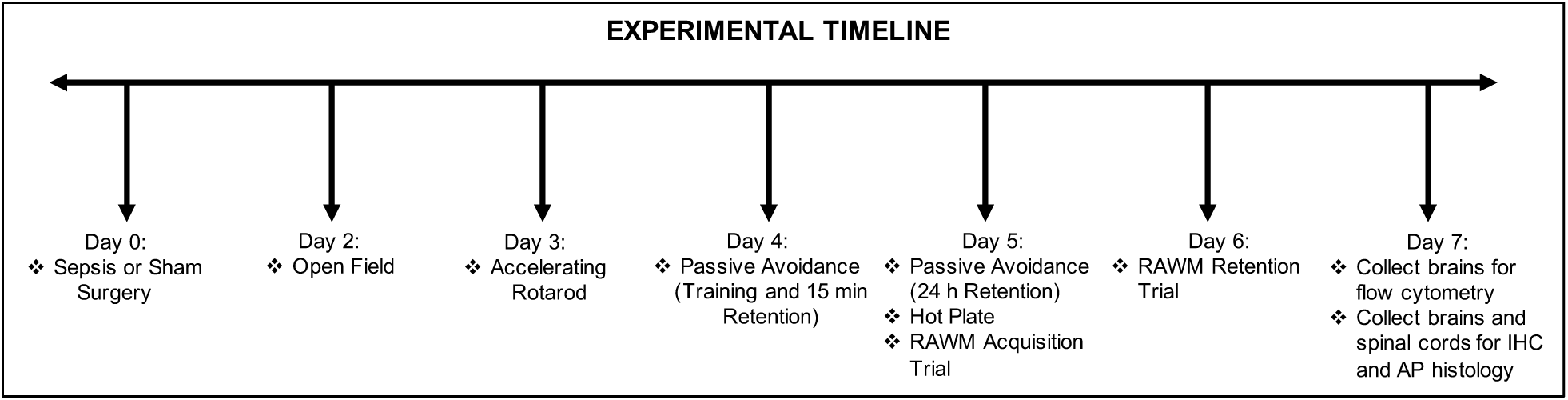
Experimental design and behavioral testing paradigm to study neuroimmune and brain microvascular function in late sepsis. Animals were randomized and underwent either sham or sepsis (cecal ligation and puncture, CLP) surgery. All animals were subjected to a series of daily behavioral assays from day 2 to day 6. Mice were euthanized on day 7 and their brains were harvested for flow cytometry, immunohistochemistry (IHC), and alkaline phosphatase (AP) histology. Spinal cords were used for IHC and AP histology.

## 3. Results

### 3.1. Survival analysis, sickness score, and average weight loss

First, we investigated the detrimental effects of sepsis on mortality and sickness behavior. Septic mice had a significantly increased mortality rate compared to sham mice (\*\*\**p* = 0.001, Log-rank) (Fig. 2A). Next, we examined the sickness score and average weight loss using the modified murine sepsis score (MMSS). Our results showed a significantly increased sickness score in septic mice compared to sham mice (F (1, 26) = 22.55, \*\*\*\**p* < 0.0001, repeated measures two-way ANOVA) (Fig. 2B). Furthermore, septic mice had a significantly higher average weight loss score compared to sham mice (U (104, 302) = 26, \*\*\**p* = 0.0005, Mann-Whitney test) (Fig 2C). Due to the downward trend in the sickness score and an improvement in overall physical appearance of septic animals after day 5, we examined blood serum levels of inflammatory cytokines interleukin 6 (IL-6) and monocyte chemoattractant protein-1 (MCP-1) chemokine on day 7 in a terminal blood sample. Our results show that septic mice exhibit a persistent peripheral inflammatory phenotype denoted by the significant increase in the levels of inflammatory cytokines IL-6 (U (16, 62) = 1, \*\**p* = 0.005, Mann-Whitney test) and MCP-1 (U (11, 55) = 1, \**p* = 0.01, Mann-Whitney test) (Fig. 2D).

**Figure 2.**
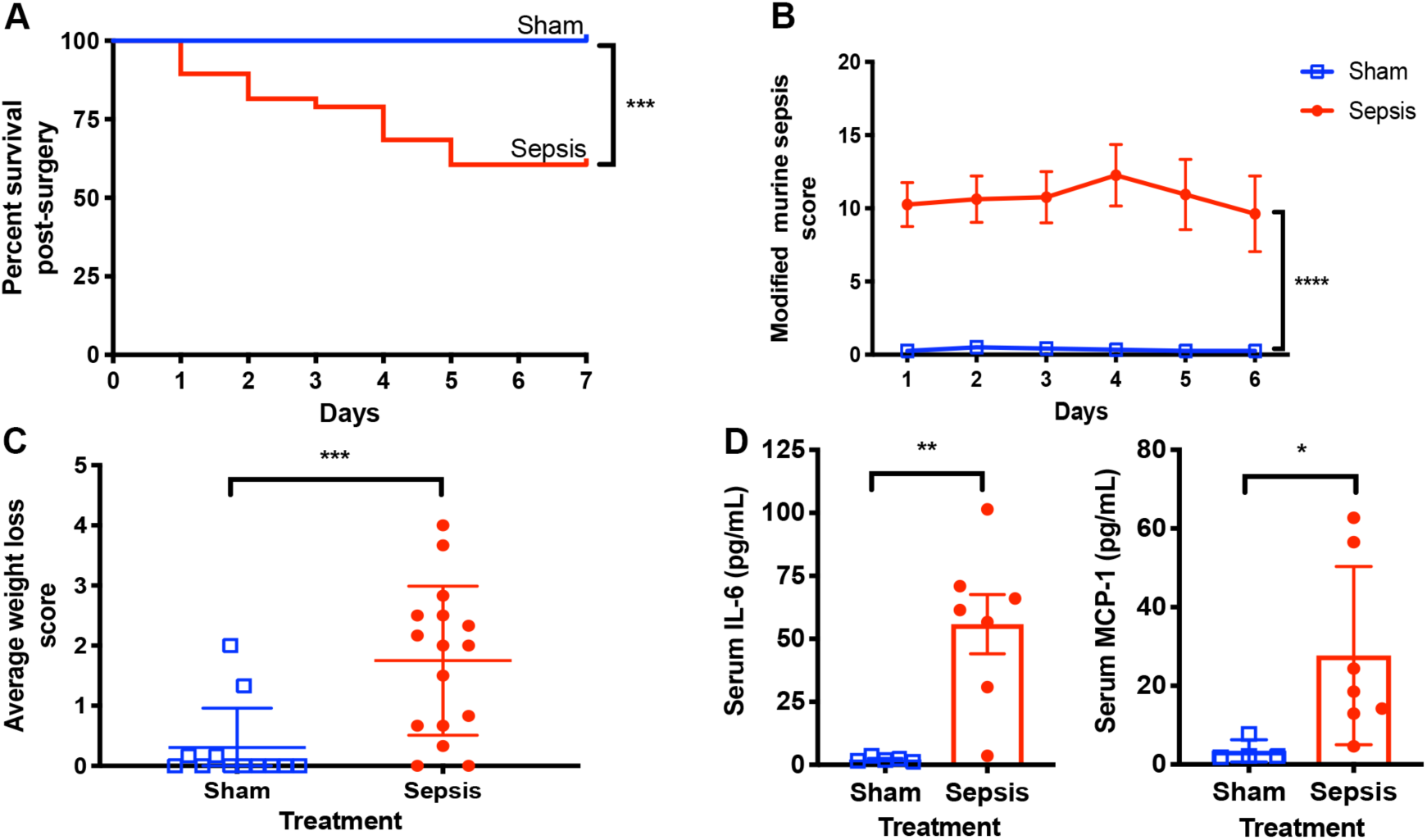
Survival is decreased while longitudinal clinical scores and serum proinflammatory cytokines are increased in sepsis. (A) A Kaplan-Meier log-rank survival curve analysis of sham (n = 22) and septic (n = 38) mice from day 0 of surgery to euthanasia at day 7 showed that septic mice had a significant decrease in survival (\*\*\**p* = 0.001) compared to sham mice. (B, C) Septic mice (n = 16) had significantly higher sickness scores (\*\*\*\**p* < 0.0001, repeated two-way ANOVA) and average weight loss scores (\*\*\**p* = 0.0005, Mann-Whitney test) compared to sham mice (n = 12). (D) Examination of blood serum inflammatory cytokines at day 7 revealed a persistent increase in IL-6 (\*\**p* = 0.005, Mann-Whitney test) and MCP-1 (\**p* = 0.01, Mann-Whitney test) in septic mice (n=7) compared to sham mice (n = 4-5). * indicates *p* < 0.05, \*\**p* <0.01, \*\*\**p* < 0.001, \*\*\*\**p* < 0.0001, and is considered significant. All data are presented as mean ± SEM.

### 3.2. Sepsis alters critical components that maintain brain microvascular barrier function

As reported previously, the expression of the tight junction (TJ) protein claudin-5 is decreased at 24 h following either CLP or LPS injection (Banks et al., 2015; Yang et al., 2015). We hypothesized that the persistence of systemic inflammatory cytokines IL-6 and MCP-1 (Fig. 2D) at day 7 in late sepsis would sustain the loss of brain microvascular integrity beyond 24 h. To investigate this, we co-immunolabelled for the endothelial cell marker CD31 (Nwafor et al.) and claudin-5 (red) in the somatosensory cortex. Our results showed that claudin-5 expression around CD31-labelled microvasculature is decreased in septic mice compared to sham mice (Fig. 3A). To further demonstrate the loss of brain microvascular integrity in sepsis we also quantified *in vivo* AP activity since a previous study reported that TNAP preserves brain endothelial integrity *in vitro* (Deracinois et al., 2015). It is important to note that TNAP is the only AP isoform found in microvessels of the brain and spinal cord (Brun-Heath et al., 2011; Street et al., 2013). Therefore, assessment of total AP activity is adequate to assess total TNAP activity. Our *in vivo* results demonstrated a significant decrease in TNAP enzyme activity in the somatosensory cortex (t = 2.8, *\*p* = 0.02, unpaired t-test) (Fig. 3B, C) of septic mice. Parallel AP histology of the spinal cord also showed decreased TNAP enzyme activity in the spinal cords of septic mice **(Supplementary Fig. 1)**. Co-labeling of TNAP enzyme activity (blue) with cerebral microvessels (CD31, brown) provides further support that the observed decrease in TNAP’s enzyme activity is not the result of a loss of cerebral microvessels, and thus most likely demonstrates that TNAP enzyme activity is reduced in cortical cerebral microvessels during late sepsis (Fig. 3D). Collectively, our findings demonstrate that loss of brain microvascular integrity in the cortex is sustained up to day 7 post-sepsis. In contrast, we did not observe any significant differences in TNAP enzyme activity in the hippocampal CA1 or CA3 region of septic and sham mice on day 7 (data not shown).

**Figure 3.**
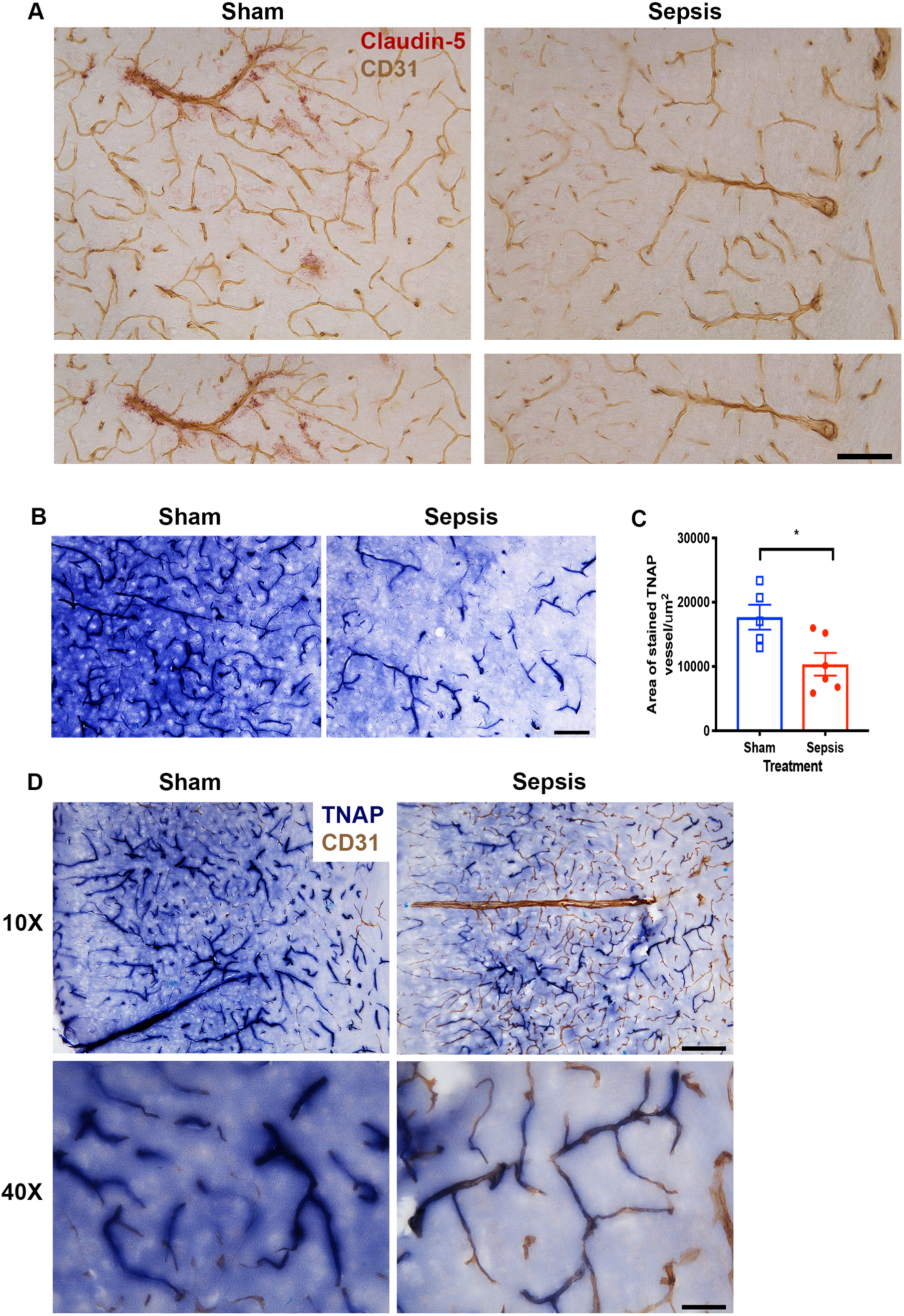
Changes in claudin-5 expression are consistent with diminished tissue nonspecific alkaline phosphatase (TNAP) enzyme activity in cerebral microvessels. (A) Claudin-5 (red) expression surrounding CD31 (brown) labelled brain microvessels is decreased in septic compared to sham mice (20X magnification). (B) Representative images of TNAP enzyme activity on brain microvessels showed that TNAP’s enzyme activity in the somatosensory cortex (20X magnification) of septic mice is decreased compared to sham mice. (C) Quantification of TNAP enzyme activity on sections of the somatosensory cortex showed that TNAP enzyme activity is significantly decreased (*\*p* = 0.02, unpaired t-test) in septic (n = 6) compared to sham treated mice (n = 5). (D) Co-labelling of TNAP enzyme activity (blue) with CD31 cerebral microvessel marker (brown) in the somatosensory cortex (10X and 40X magnification) further demonstrates that loss of TNAP’s enzyme activity in brain sections is due to a loss of TNAP enzyme activity in cerebral microvessels rather than a loss of brain microvessels. * indicates *p* < 0.05, and is considered significant. All data are presented as mean ± SEM. 10X, 20X, and 40X magnification scale bar = 160 µm, 80 µm, and 40 µm respectively.

### 3.3. Dysfunction of the neuroimmune axis post-sepsis

#### 3.3.1. Sustained peripheral myeloid immune cell trafficking into brain parenchyma of septic mice

To investigate whether alterations in TNAP activity within cerebral microvessels (Fig. 3A, B) are coupled to trafficking of peripheral immune cells into the brain parenchyma at day 7 post-sepsis, we next characterized the percentage of myeloid cell populations in the brains of septic and sham mice using flow cytometry. Myeloid cells are gated using the strategy shown in **Supplementary Figure 2**. FCS express analyses of inflammatory myeloid cell populations showed that neutrophils (CD45^hi^, CD11b^+^, Ly6C^-^, Ly6G^+^) (Fig. 4A, B) and monocytes (CD45^hi^, CD11c^+^, Ly6C^+^, Ly6G^-^) (Fig. 4C, D) are significantly increased (U (15, 51) = 0, \*\**p* = 0.004, Mann-Whitney test) in the brains of septic compared to sham mice. Anti-inflammatory myeloid derived suppressors cells (MDSCs) i.e. polymorphonuclear-MDSCs/PMN-MDCs (CD45^hi^, CD11b^+^, Ly6C^lo^, Ly6G^+^) (Fig. 4E, F; U (15, 51) = 0, \*\**p* = 0.004, Mann-Whitney test) and monocytic-MDSCs/M-MDSCs (CD45^hi^, CD11b^+^, Ly6C^hi^, Ly6G^-^) (Fig. 4E, G; U (15, 51) = 0, \*\**p* = 0.004, Mann-Whitney test) were also significantly increased in the brains of septic compared to sham mice.

**Figure 4.**
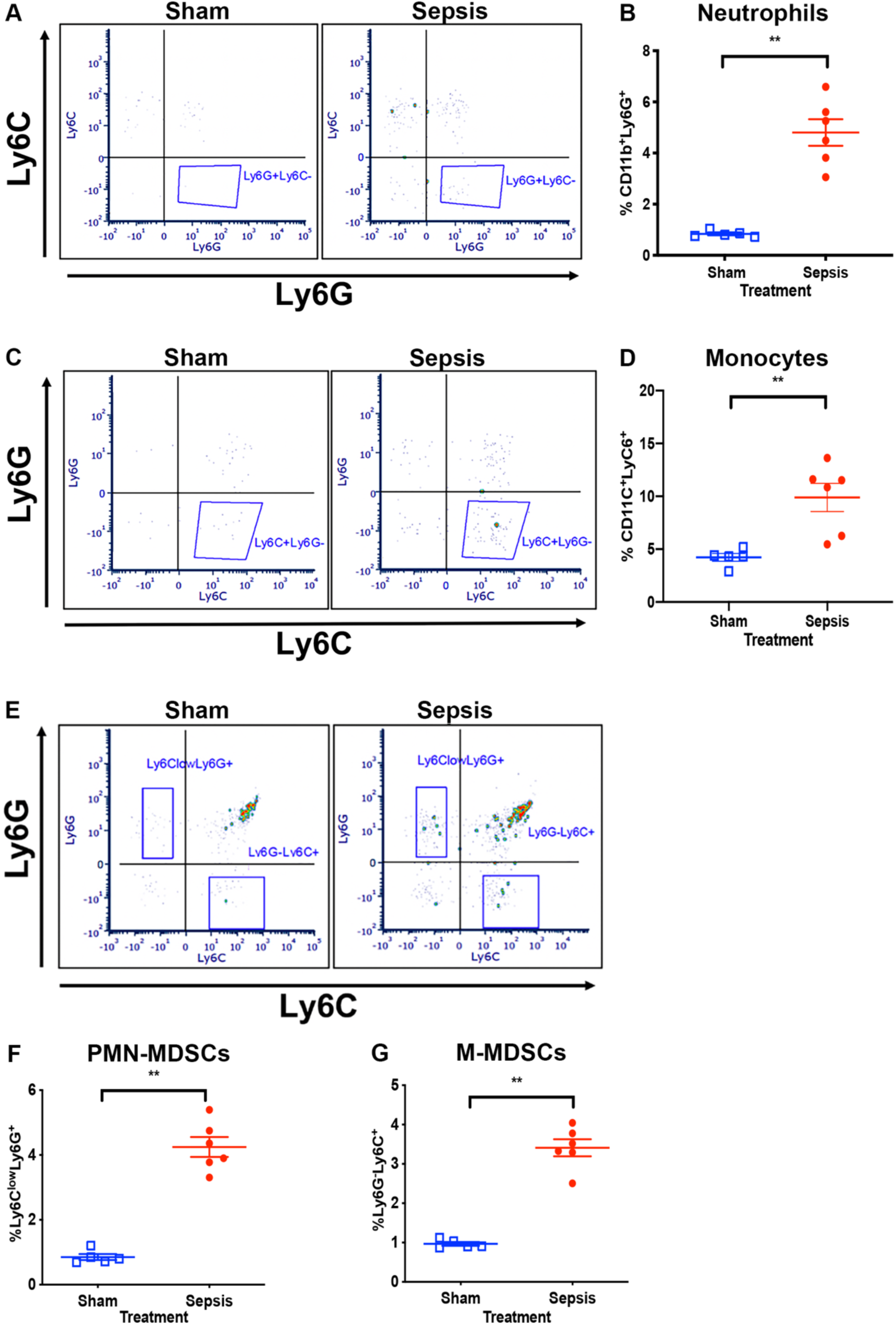
Brain infiltrating myeloid cell populations are elevated in late sepsis. (A-D) Representative dot plot and quantification of inflammatory myeloid cells showed a significant increase in neutrophil (\*\**p* = 0.004, Mann-Whitney test) and monocyte (\*\**p* = 0.004, Mann-Whitney test) cell populations in the brains of septic compared to sham mice. (E-G) Analyses of anti-inflammatory myeloid derived suppressor cells (MDSCs) showed that septic brains have a significant increase in polymorphonuclear-MDSC/PMN-MDSC (\*\**p* = 0.004, Mann-Whitney test) and monocytic-MDSCs/M-MDSC (\*\**p* = 0.004, Mann-Whitney test) cell populations compared to sham mice. Flow cytometric results are expressed as the percentage of an indicated cell type in total CD45^+^ live cells in septic mice (n = 6) or sham mice (n = 5). Analyzed population is indicated by the encircled gating. ** indicates *p* < 0.01, and is considered significant. All data are presented as mean ± SEM.

#### 3.3.2. Sustained peripheral T-lymphoid immune cell trafficking into brain parenchyma of septic mice

Next, we explored T-lymphoid immune cell trafficking post-sepsis using the gating strategy for CD45^hi^ live cells shown in **Supplementary Figure 2**. FCS express analyses of CD8+ (CD45^hi^, CD3^+^, CD8^+^) (Fig. 5A, B; U (15, 51) = 0, \*\**p* = 0.004, Mann-Whitney test) and CD4+ T-cells (CD45^hi^, CD3^+^,CD4^+^) (Fig. 5C, D; t = 2.3, *\*p* = 0.046, unpaired t-test) revealed increased populations of these cells in the brains of septic compared to sham mice. Owing to the observed increase in anti-inflammatory MDSCs post-sepsis (Fig. 4E-G), we assessed whether anti-inflammatory cells (T-regulatory cells) from the T-lymphoid lineage were similarly increased in the brains of septic mice. To do this, CD4+ T-cells are gated further for Foxp3 and CD25 markers. Our results showed that T-regulatory cells (CD4^+^, CD25^+^, Foxp3^+^) (Fig. 5E, F) are significantly increased (t = 4.7, *\*\*p* = 0.0011, unpaired t-test) in the brains of septic mice compared to sham mice. Together both findings from Figures 4A-G and 5A-F demonstrate a persistent dysfunction in the neuroimmune axis in septic mice.

**Figure 5.**
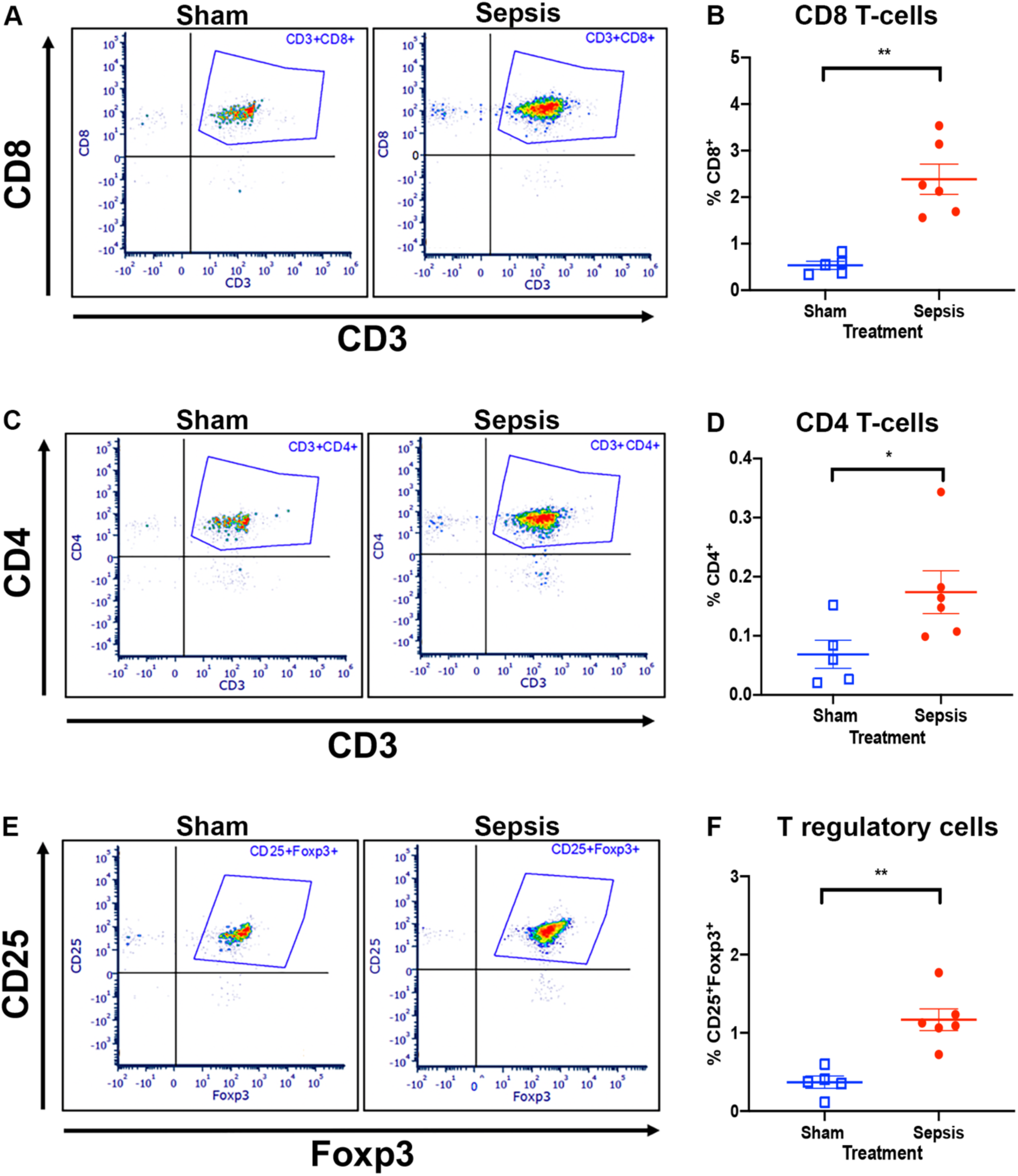
Multiple T-lymphoid cell populations are elevated in the brain during late sepsis. (A-D) Representative dot plot and quantification of T-lymphoid cells revealed that septic brains have a significant increase in CD8+ (\*\**p* = 0.004, Mann-Whitney test) and CD4+ T-cell (*\*p* = 0.046, unpaired t-test) populations compared to sham mice. (E, F) Further analyses of a specific subset of CD4^+^ cells (T-regulatory cells, Tregs) using Foxp3 and CD25 markers revealed a significant increase in the percentage of Tregs (*\*\*p* = 0.0011, unpaired t-test) in the brains of septic compared to sham mice. Quantification of flow cytometric results are expressed as the percentage of an indicated cell type in total CD45^+^ live cells in septic mice (n = 6) or sham mice (n = 5). The analyzed population is indicated by the encircled gating. * indicates *p* < 0.05, \*\**p* < 0.01, and is considered significant. All data are presented as mean ± SEM.

### 3.4. Whole brain and regional analyses of brain residential microglia populations in late sepsis

Our next step was to determine the effects of sepsis on brain residential microglia cell populations. For our initial analysis, we used flow cytometry to evaluate whole brain residential microglia populations post-sepsis using the gating strategy CD45^low^, CD11b^+^, and CD11c^-^ (**Supplementary Figure 3 and** **Fig. 6A**). Interestingly, our flow cytometric analyses showed that septic mice had a significant decrease in whole brain residential microglia cell populations (CD45^low^, CD11b^+^, CD11c) (U (45, 21) = 0, \*\**p* = 0.004, Mann-Whitney test) compared to sham mice (Fig. 6A, B). To ascertain whether these results were comparable to region-specific changes in microglial abundance, we performed immunohistochemistry and assessed microglia in three different brain regions (i.e. somatosensory cortex, CA1, and CA3) for the microglia/macrophage ionized calcium binding adaptor molecule 1 (Iba-1). Contrary to the results shown in our flow cytometric whole brain analyses (Fig. 6A, B), Iba-1 positively stained cells were significantly increased in the somatosensory cortex (U (15, 51) = 0, \*\**p* = 0.004, Mann-Whitney test), and CA1 (t = 3.8, *\*\*p* = 0.004, unpaired t-test) and CA3 (t = 3.6, *\*\*p* = 0.006, unpaired t-test) hippocampal regions of the brain (Fig. 6C-H). To confirm whether the stained Iba-1 positive cells were microglia and not macrophages, we stained for microglia using a specific marker, (TMEM119), which is not found on macrophages (Bennett et al., 2016). Contrary to the observed increase in Iba-1 positive cells shown in Figures 5 C-H, there were no significant differences in TMEM119 positive cells in the somatosensory cortex (t = 0.22, *p* = 0.13, unpaired t-test), and the CA1 (t = 0.07, *p* = 0.94, unpaired t-test) and CA3 (t = 0.42, *p* = 0.68, unpaired t-test) hippocampal regions **(Supplementary Table 1)**.

**Figure 6.**
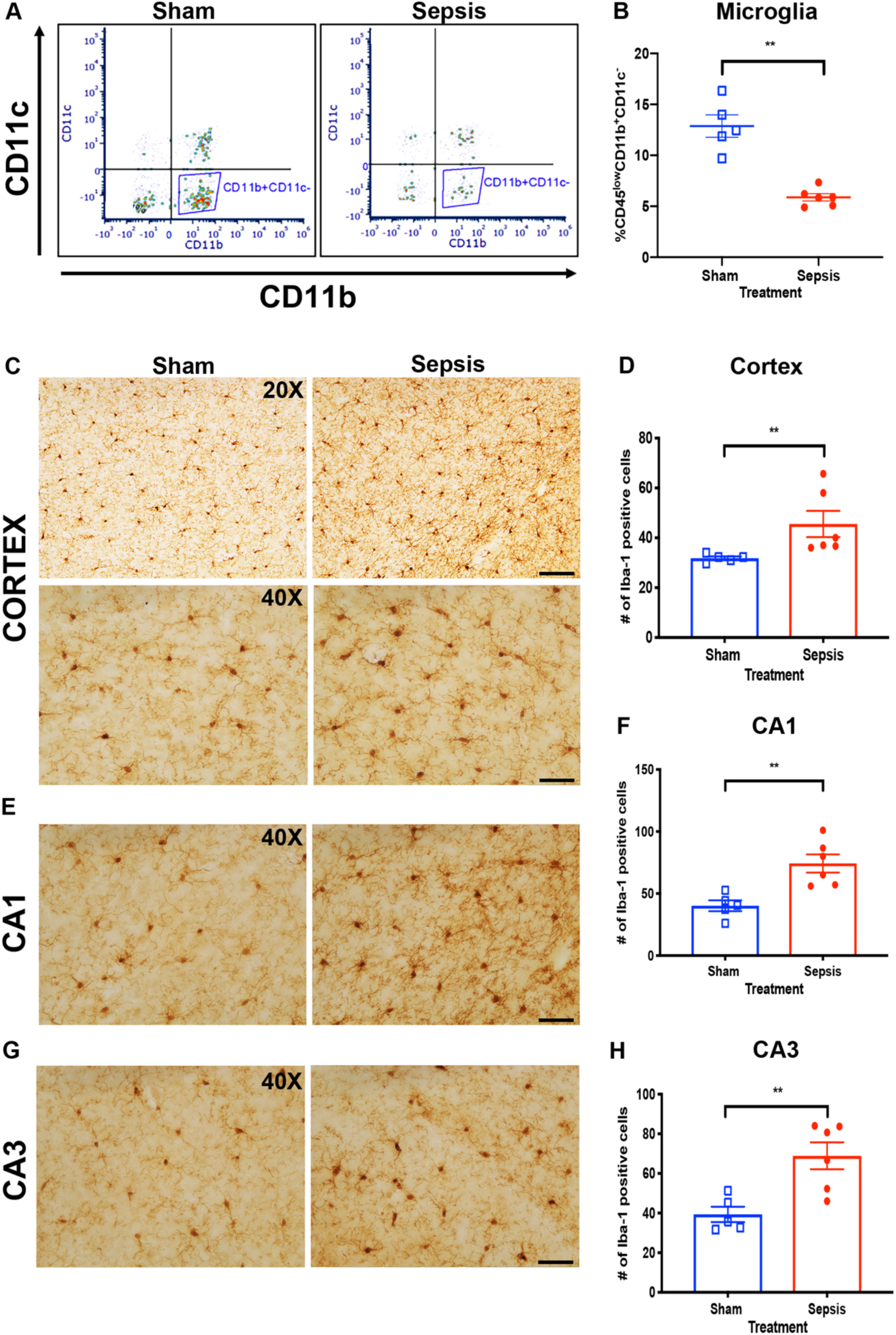
Late sepsis is characterized by heterogeneity in microglia/infiltrating monocyte populations. (A, B) Representative dot plot and quantification of whole brain residential microglia population by flow cytometry revealed a significant decrease in the percentage of microglia (\*\**p* = 0.004, Mann-Whitney test) in the brains of septic compared to sham mice. Quantification of flow cytometric results is expressed as the percentage of microglia in total CD45^+^ live cells in septic mice (n = 6) or sham mice (n = 5). Analyzed population is indicated by the encircled gating. (C-F) Representative histological images (C,E,G) and quantification of Iba-1 positive cells (D,F,H) showed an increased number of Iba-1 positive cells in the somatosensory cortex (\*\**p* = 0.004, Mann-Whitney test), CA1 (*\*\*p* = 0.004, unpaired t-test) and CA3 (*\*\*p* = 0.006, unpaired t-test) brain regions of septic (n = 6) compared to sham mice (n = 5). ** indicates *p* < 0.01, and is considered significant. All data are presented as mean ± SEM. 20X and 40X magnification scale bar = 80 µm, and 40 µm respectively.

### 3.5. Sustained astrogliosis in the somatosensory cortex of septic mice

Given the alterations in brain microglial/monocyte activation, we also assessed changes in astrocyte proliferation and morphology post-sepsis. We performed glial fibrillary acidic protein (GFAP) immunolabeling and quantified the proliferation of reactive astrocytes in the somatosensory cortex, in hippocampal CA1 and CA3 and in the spinal cord. Our results demonstrate a significant increase in astrocyte proliferation in the somatosensory cortex (t = 3.5, *\*\*p* = 0.006, unpaired t-test) of septic compared to sham mice (Fig. 7A, B). However, we did not observe any significant differences in astrocyte proliferation in the CA1 (t = 0.5, *p* = 0.61, unpaired t-test) and CA3 (t = 0.60, *p* = 0.56, unpaired t-test) hippocampal regions of the brain (Fig. 7C-F). Parallel to the results observed in the somatosensory cortex of the brain, we observed an increased number of GFAP positive cells and increased proliferative astrocytes in the spinal cords of septic mice compared to sham mice **(Supplementary Figure 4A, B)**. These results demonstrate that the somatosensory cortex and the spinal cord exhibit sustained astrogliosis post-sepsis.

**Figure 7.**
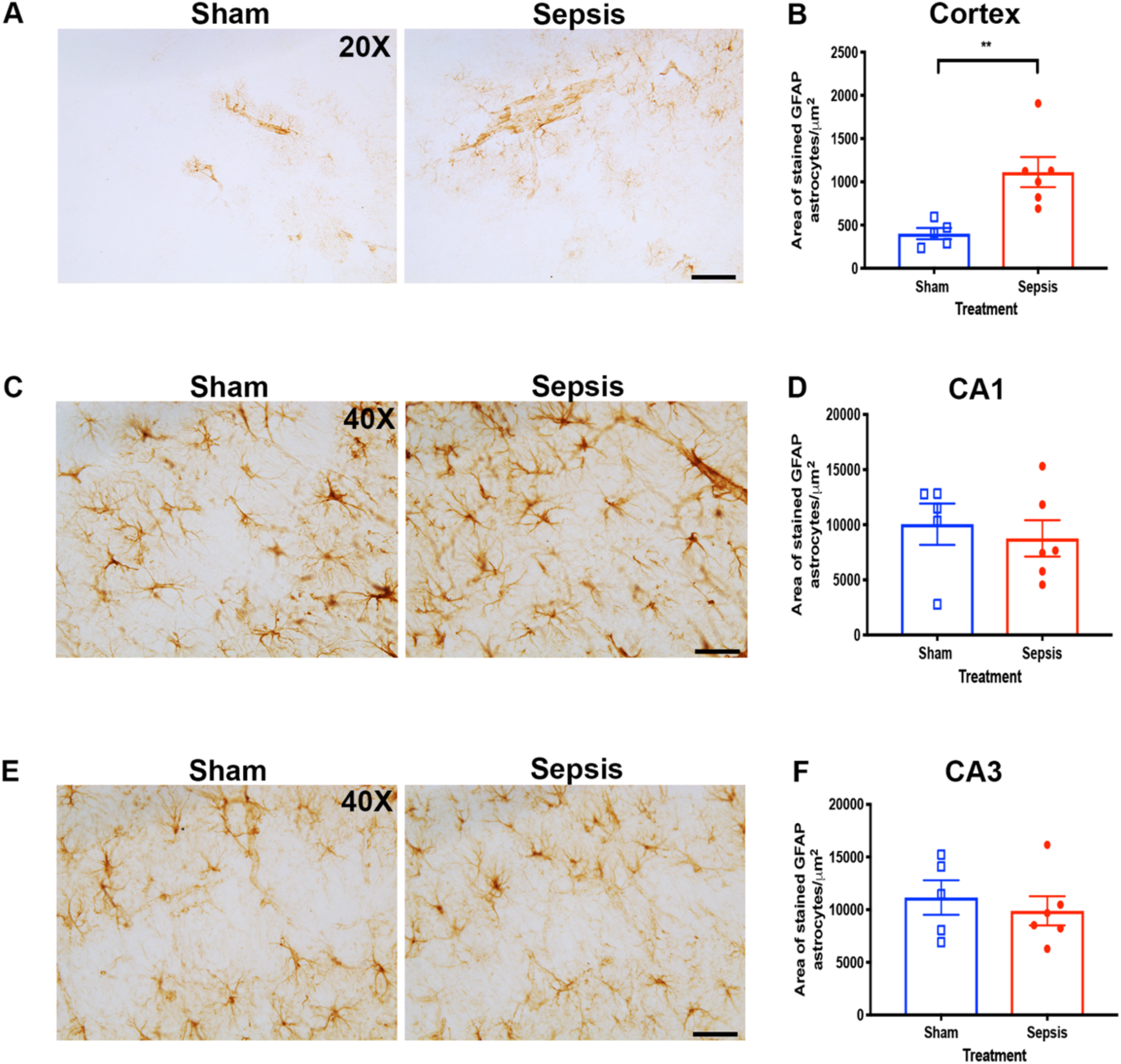
Sustained astrogliosis in the somatosensory cortex of septic mice. (A-F) Representative glial fibrillary acidic protein (GFAP) astrocyte histological images and quantification of astrogliosis (astrocyte proliferation) in the somatosensory cortex, CA1, and CA3 brain regions of septic and sham mice. (B) Quantification of GFAP astrocytes showed a sustained significant increase in astrocyte proliferation in the somatosensory cortex (*\*\*p* = 0.006, unpaired t-test) of septic (n = 6) compared to sham mice (n = 5). (D, F) However, no differences in astrocyte proliferation were seen between septic (n = 6) and sham mice (n = 6) in the hippocampal brain regions CA1 (*p* = 0.61, unpaired t-test) and CA3 (*p* = 0.56, unpaired t-test). ** indicates *p* < 0.01, and is considered significant. All data are presented as mean ± SEM. 20X and 40X magnification scale bar = 80 µm, and 40 µm respectively.

### 3.6. Septic mice displayed an impairment in spontaneous locomotion and exhibited a novel anti-nociception phenotype

Septic and critically ill patients often experience long-term sensorimotor impairment (Iwashyna et al., 2010; Khan, Harrison, Rich, & Moss, 2006; Zauner et al., 2002). Therefore, we implemented a series of behavioral assays on days 2 (open field testing, OFT) and 3 (rotarod) to assess motor dysfunction and on day 5 (hot plate) to assess sensory dysfunction in septic mice. The OFT was used to evaluate spontaneous motor dysfunction while the rotarod was utilized to assess evoked motor dysfunction. Our results showed that spontaneous locomotion in the horizontal (F (1, 24) = 20.51, \*\*\**p* < 0.0001, repeated measures two-way ANOVA) and vertical axes (F (1, 24) = 48.4, \*\*\**p* < 0.0001, repeated measures two-way ANOVA) were significantly decreased in septic compared to sham mice on day 2 over a 60 min trial (Fig. 8A, B). However, we did not observe any differences in evoked locomotion on the rotarod (F (1, 13) = 1.6, *p* = 0.23, repeated measures two-way ANOVA) between septic and sham mice over four 300 s trials (Fig. 8C). Next, we utilized the hot plate test to assess sensory dysfunction post-sepsis. Our results show that septic mice have an increased latency to respond to thermal stimuli (t = 2.8, *\*\*p* = 0.01, unpaired t-test) and exhibit a decreased total number of nociceptive behaviors (hind limb-lick, flick, and jump) (t = 2.9, *\*\*p* = 0.0096, unpaired t-test) compared to sham mice (Fig. 8D, E). From these findings, we report, for the first time, a discrepancy between two measurements of locomotion, OFT and rotarod, in septic mice, and reveal a novel anti-nociceptive phenotype exhibited by septic mice.

**Figure 8.**
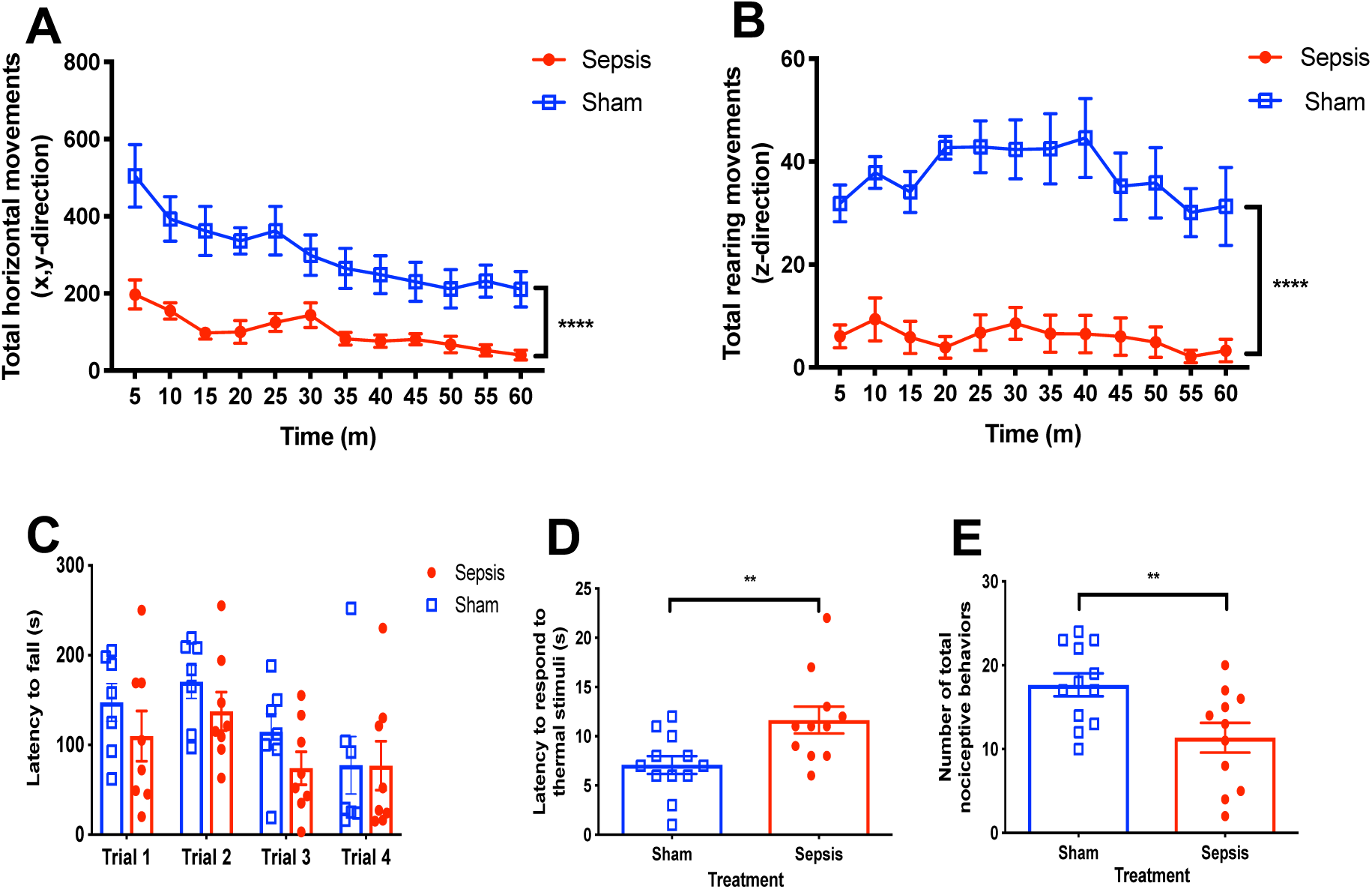
Spontaneous sensorimotor activity and nociceptive functions are impaired in septic mice. (A, B) Open field testing of spontaneous locomotion on day 2 showed that septic mice (n = 14) had a significant decrease in horizontal (***p < 0.0001, repeated two-way ANOVA) and vertical (***p < 0.0001, repeated two-way ANOVA) spontaneous locomotion compared to sham mice (n = 10). (C) Rotarod assessment of evoked locomotion on day 3 showed no difference (*p* = 0.23, repeated two-way ANOVA) between septic mice (n = 8) and sham mice (n = 7). (D, E) Evaluation of sensory dysfunction with the hot plate test revealed a novel antinociceptive behavior in septic mice i.e. septic mice exhibited a significant increase in the latency to respond to thermal stimuli (*\*\*p* = 0.01, unpaired t-test) and a significant decrease in the total number of nociceptive behaviors (hind limb - lick, flick, and jump) (*\*\*p* = 0.0096, unpaired t-test). ** indicates *p* < 0.01, \*\*\*\**p* < 0.0001, and is considered significant. All data are presented as mean ± SEM.

### 3.7. Learning and memory is preserved seven days post-sepsis

To determine whether learning and memory deficits are evident during late sepsis, we used the two-day radial arm water maze (2D-RAWM) to assess for spatial learning and memory on days 5 and 6, and the passive avoidance test to assess non-spatial learning and memory on days 4 and 5. Septic mice showed no difference in the number of errors (F (1, 20) = 0.76, *p* =0.39, repeated two-way ANOVA) or the latency to find the hidden platform (F (1, 20) = 0.87, *p* = 0.36, repeated two-way ANOVA) on day 2 of the retention trial compared to sham mice (Fig. 9A, B). Assessment of non-spatial learning and memory with the use of the passive avoidance test showed no difference in latency to enter the darkened compartment (F (1, 12) = 0.09, *p* = 0.77, repeated two-way ANOVA) between septic and sham mice at the 15 min and 24 h retention trials (Fig. 9C). Due to the absence of learning and memory impairments at these timepoints, we assessed neuronal loss via NeuN immunolabeling in hippocampal regions CA1 and CA3. No apparent differences in neuronal loss were observed in these regions between sham and septic mice **(Supplementary Figure 5)**. To confirm that other brain regions involved in memory consolidation were not affected by sepsis, we performed immunostaining for choline acetyltransferase (ChAT) in cholinergic neurons of the basal forebrain. Similarly to results observed in the hippocampus, there were no significant differences in the number of ChAT positive neurons in the basal forebrain of septic mice compared to sham mice **(Supplementary Table 2)**. Taken together, the absence of neuronal loss in the hippocampus and cholinergic neurons of the basal forebrain in septic mice further substantiated the results derived from the 2D-RAWM and passive avoidance behavioral tests (Fig. 9A-C).

**Figure 9.**
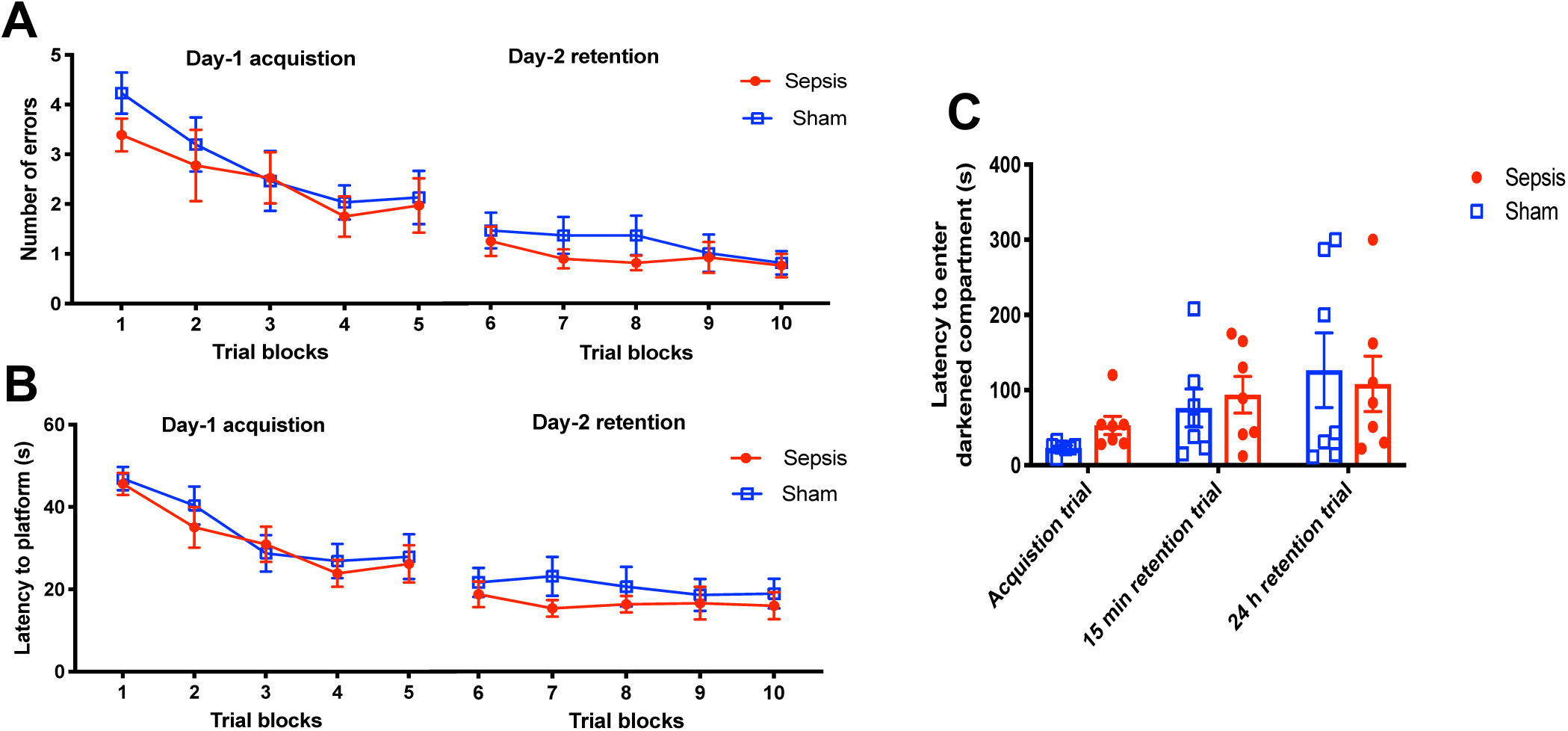
Absence of spatial and non-spatial learning and memory behavioral deficits in late sepsis. (A, B) Mice underwent two-day radial arm water maze (2D-RAWM) testing to assess spatial learning and memory deficits on days 5 and 6 post-sepsis. Septic mice (n = 12) showed no difference in the number of errors (*p* = 0.39, repeated two-way ANOVA) or the latency to find the hidden platform (*p* = 0.36, repeated two-way ANOVA) on day 2 of the retention trial compared to sham mice (n = 10). (C) The passive avoidance assessed non-spatial learning and memory on days 4 and 5 post-sepsis. Septic mice (n = 7) showed no difference in latency to enter the darkened compartment (*p* = 0.77, repeated two-way ANOVA) compared to sham mice (n = 7). All data are presented as mean ± SEM.

## 4. Discussion

In this study, we employed the cecal ligation and puncture model of experimental sepsis to identify neuroimmune mechanisms and behavioral deficits that coupled a potential novel *in vivo* marker of brain microvascular dysfunction in sepsis, i.e. the loss of TNAP activity in cerebral microvessels. Brain function in humans is significantly impaired in both early and late sepsis and is an independent predictor of mortality (Gofton & Young, 2012). In sepsis patients, slow delta activity on an electroencephalogram (EEG) is associated with severe neurocognitive decline (Hosokawa et al., 2014). More importantly, the detection of peripherally injected gadolinium and the accumulation of vasogenic fluid in the brains of septic rodents are evidence for brain microvascular dysfunction during sepsis (Bozza et al., 2010; Towner et al., 2018). An emerging hypothesis suggests that the pronounced hyper-inflammatory phase seen in early sepsis disrupts brain microvascular function and barrier integrity (Mazeraud et al., 2016; Widmann & Heneka, 2014). The subsequent infiltration of peripheral immune cells and systemic pro-inflammatory molecules into the brain parenchyma that accompanies changes in brain microvascular dysfunction may increase the likelihood of significant short-term and long-term deficits in neurocognitive function (Andonegui et al., 2018; Sonneville et al., 2013). Therefore, a better understanding of brain microvascular mechanisms that maintain barrier integrity may foster the development of therapeutics to mitigate the neurological impairments seen in sepsis survivors. Our results suggest that TNAP is a plausible regulator of brain microvascular function in experimental sepsis, and potentially, other neurological disorders with cerebrovascular dysfunction.

First, we validated our sepsis model by evaluating mortality, sickness behavior, and weight loss post-sepsis. Septic mice exhibited increased mortality, sickness behavior, and average weight loss compared to sham mice. These results align with previously reported findings in similar models of sepsis (Bi et al., 2019; Zaghloul et al., 2017). Although the sickness score began to improve after day 4 and no further deaths were recorded after day 5, overall clinical deficits were contrary to the levels of circulating inflammatory cytokines IL-6 and MCP-1, which were significantly increased in septic mice. Intravenous injection of pro-inflammatory cytokines or treatment of mouse brain capillary endothelial cells with pro-inflammatory cytokines have been shown to promote brain microvascular dysfunction (Blamire et al., 2000; Nishioku et al., 2010; Varatharaj & Galea, 2017). Therefore, we performed immunohistochemistry to assess the spatial localization of claudin-5, a well-characterized tight junction (TJ) protein found on brain endothelial cells, to investigate alterations in brain microvascular integrity. Our results showed that claudin-5 expression is decreased around the CD31 positive cerebral microvessels in septic mice. Next, we examined whether an increase or decrease in TNAP activity was concomitant with the loss of claudin-5 expression post-sepsis. Our findings showed that TNAP enzyme activity is significantly decreased on CD31 positive cerebral microvessels in late sepsis. This finding corroborates and expands on our previously published data showing that TNAP’s activity is decreased in septic mice 24 h post-sepsis (Nwafor et al., 2019). However, the observed decrease in TNAP activity at day 7 post-sepsis was localized to somatosensory cortex and spinal cord and absent from the hippocampus.

Our next objective was to identify alterations in brain myeloid and lymphoid cell populations in order to elucidate the patterns of immune cell trafficking which parallel the loss of brain microvascular integrity. While alterations in peripheral myeloid and T-lymphoid cell populations have been described in numerous murine sepsis studies (Deutschman & Tracey, 2014; Hotchkiss, Coopersmith, McDunn, & Ferguson, 2009), few studies have addressed changes in brain immune cells populations and trafficking peripheral leukocyte populations in late sepsis. Our results showed a persistent dysfunction of the neuroimmune axis at day 7 post-sepsis as demonstrated by the increase in brain myeloid and T-lymphoid populations in the brain of septic mice compared to sham mice. This result is consistent with findings from Singer *et al*. who also showed that monocytes and neutrophils persist up to day 14 in septic rodent brains (Singer et al., 2016). More importantly, our finding emphasizes that elevated numbers of leukocytes are present in the brain parenchyma post-sepsis in spite of a downward trend in sickness score, physical appearance of animal, and cessation of additional mortality past day 5. Due to the persistent dysfunction of the neuroimmune axis post-sepsis, we investigated the impact of these changes on brain microglial populations. Recent findings suggest that brain microglial depletion exacerbates brain and systemic inflammation and that repopulating microglia enhances neurological recovery post-sepsis (Michels et al., 2019). Flow cytometric analysis of whole brain microglial populations (CD45^low^, CD11b^+^, CD11c^-^) post-sepsis showed that these cells are decreased, which suggests that the loss of the resident microglia cells is a hallmark associated with post-sepsis neurological impairment. However, further studies are needed to delineate whether the beneficial effects of brain microglial cell repopulation post-sepsis as demonstrated by Michels *et al*. (Michels et al., 2019) are associated with the immunophenotype CD45^low^, CD11b^+^, and CD11c^-^ found in our study or another predominant microglia phenotype.

Additionally, we also uncovered two novel findings that support a role for brain immunosuppression in late sepsis. It has been demonstrated that sepsis-associated immunosuppression in peripheral tissues contributes to poor recovery and long-term morbidity post-sepsis (Delano & Ward, 2016; Mira, Brakenridge, Moldawer, & Moore, 2017), but a role of brain immunosuppression in sepsis is less clear. First, we observed that brain CD4^+^/CD25^+^/Foxp3^+^ T-regulatory cells (Tregs) were elevated in septis mice. Although dysregulation of peripheral Tregs may play a role in generating the immunosuppressive phenotype associated with late sepsis (Cao et al., 2018; Molinaro et al., 2015; Tatura et al., 2015), there are no reports, to our knowledge, that have investigated brain Treg populations in sepsis. The elevation in brain Treg populations we identified in septic brains suggests a putative role for these cells in brain-mediated immunosuppression; however, further studies are needed to substantiate this finding. Second, we assessed whether myeloid derived suppressor cells (MDSCs), a peripheral immune cell population with T-cell suppressive capacity, were also present in brain tissue. We found that both PMN-MDSCs and M-MDSCs were elevated at seven days post sepsis. While the role of MDSCs in neuroinflammation is not well understood, a recent study in a mouse model of traumatic brain injury showed that circulating MDSCs entered the brain shortly after injury to suppress neuroinflammation (Hosomi et al., 2019). In contrast, the results from our study suggest that the sustained presence of both PMN-MDSCs and M-MDSCs in late sepsis supports an immunosuppressive role associated with poor outcome. From our results, we speculate that Tregs and MDSCs play a critical role in post-sepsis immunosuppression in the brain, which, in turn, may contribute to long-term, post-sepsis neurological impairment.

Regional analyses of microglia populations using Iba-1 and CD11b as markers have shown that microglial numbers are increased in the cortex and hippocampus of septic mice (Hoogland, Houbolt, van Westerloo, van Gool, & van de Beek, 2015; Singer et al., 2016). We used the Iba-1 to quantify microglia/infiltrating monocyte populations in the somatosensory cortex and hippocampus (CA1 and CA3 regions). In agreement with prior reports, we found that Iba1 positive cell populations were increased in cortex and hippocampus. However, it is important to note that the Iba-1 and CD11b markers label both microglia and infiltrating monocytes (Imai, Ibata, Ito, Ohsawa, & Kohsaka, 1996). To discern a clearer role for residential brain microglia and infiltrating monocytes during late sepsis, further immunostaining with TMEM119, a marker for residential microglia, did not reveal any differences between microglial populations in sham and septic mice. The observed differences reflect the heterogeneity of the brain microglial/monocyte populations and emphasize their importance in the pathophysiology of sepsis (Colton, 2009; Smolders et al., 2019). Our results highlight differential outcomes pertaining to the quantification of brain residential microglia populations based on the type of analyses performed and the brain region of interest. Additionally, the resultant increase in Iba-1 positive cells we observed may be attributed to infiltrating macrophage populations from the periphery rather than an alteration in brain-resident microglial populations, which was confirmed using TMEM119. Lastly, we assessed astrogliosis in late sepsis. Our results showed that astrocyte proliferation is significantly increased in the somatosensory cortex and spinal cord of septic mice; however, no significant differences were observed in the CA1 and CA3 hippocampal brain regions between septic and sham mice. The astrogliosis results complement the significant decrease in cerebral microvessel TNAP activity in the cortex and spinal cord.

Since septic survivors are burdened with neurological impairments (Iwashyna et al., 2010), we investigated whether the neuroinflammatory findings in our sepsis model paralleled sensorimotor or cognitive impairments. First, we investigated motor dysfunction in septic animals using open field and rotarod testing. Our results showed significantly reduced spontaneous locomotion in septic compared to sham mice. However, when septic and sham mice were measured for evoked locomotion on the rotarod no differences were observed. We speculate that the significant decrease in spontaneous locomotion is due to a lack of motivation to explore a novel environment post-sepsis; however, when septic mice were forced to engage in motor activity using the rotarod test, this motor dysfunction disappears. Motivation is an affective behavior that is difficult to quantify in mice (Ward, 2016) and sepsis-induced changes in motivation have been observed following lipopolysaccharide injection (Anderson, Commins, Moynagh, & Coogan, 2015). Recent studies report similar neuropsychiatric disturbances in motivation in sepsis survivors (Barichello et al., 2019; Erbs et al., 2019).

Due to the significant loss of TNAP activity in cerebral microvessels from the somatosensory cortex and spinal cords of septic mice, we also assessed thermal sensory dysfunction using the hot plate test. Our results uncovered a novel anti-nociceptive phenotype in septic mice. Altered pain perception has previously been reported in sepsis patients (Thomas, Stover, Lambert, & Thompson, 2014), and increased sensitivity to deep pain but not cutaneous pain has been reported in studies of human experimental endotoxemia (Karshikoff et al., 2015).

We did not observe any alterations in both spatial and non-spatial learning and memory tests between septic and sham mice. When juxtaposed with our learning and memory tests, the absence of neuronal loss in the basal forebrain or the hippocampus demonstrates that learning and memory are not affected at seven days post-sepsis. It is highly likely that the seven day post-sepsis time frame was too early to detect cognitive impairment, as other studies have shown that learning and memory are significantly altered at one month post-sepsis in mice (Andonegui et al., 2018; Chavan et al., 2012).

In summary, this study establishes that diminished TNAP enzyme activity in cerebral microvessels is a consequence of the brain’s sustained response to systemic inflammation and infection, i.e. sepsis. While our results demonstrate that TNAP activity is suppressed in cortex and spinal cord, it is unclear if TNAP activity is also suppressed in other brain regions like the hippocampus at later time points post-sepsis when learning and memory deficits become more apparent. Collectively, the findings from this study provide a framework for investigating the contribution of TNAP to brain microvascular dysfunction, neuroinflammation, leukocyte trafficking, and behavioral deficits in late sepsis. A better understanding of these mechanisms will support the development of targeted therapies to mitigate chronic neurological impairments in sepsis survivors.

## Supporting information

Manuscript and Figures

## 5. Author Contributions

D.C.N, D.D, and C.M.B designed the studies. D.C.N. and A.L.B. performed CLP surgeries and monitored animals after sepsis procedure. D.C.N., C.A.G., and E.B.E-C. performed behavioral tests and analyzed behavioral data. D.C.N. and S.A.B. performed immunohistochemistry and image analysis. D.C.N., S.C., W.W., and S.J. performed and analyzed flow cytometric data. D.C.N., D.D., C.M.B. wrote the manuscript. All authors read and revised the final manuscript.

## Acknowledgements

The authors gratefully thank Jessica Povroznik, M.S. for her technical support and assistance with rodent behavioral training. We acknowledge the technical support from Dr. Kathleen Brundage in the West Virginia University Flow Cytometry and Single Cell Core Facility, which is supported by the National Institutes of Health (NIH) equipment grant number S10 OD016165 and the Institutional Development Awards (IDeA) from the National General Medical Sciences of the National Institutes of Health under grant numbers P30 GM103488 (Cancer CoBRE) and P20 GM103434 (INBRE). Funding for this work was supported by the NIH T32 AG052375 (D.C.N, A.L.B), K01 NS081014 (C.M.B), West Virginia Clinical and Translational Science Institute (U54 GM104942), and the West Virginia University Stroke CoBRE (P20 GM109098).

