## Supplementary material for "Loss of tissue non-specific alkaline phosphatase (TNAP) enzyme activity in cerebral microvessels is coupled to persistent neuroinflammation and behavioral deficits in late sepsis": Manuscript and Figures

**Supplementary Tables:**

**Supplementary Table 1**

Quantification of TMEM 119 positive brain microglia populations sham (n = 5) and septic mice (n = 6)

| Area of stained TMEM119 microglia/µm$^2$ | | | |
|---|---|---|---|
| Treatment | | | |
| Brain region | Sham (mean ± SEM) | Sepsis (mean ± SEM) | p-value |
| Cortex | 17634 ± 2313 | 27110 ± 4924 | 0.13 |
| CA1 | 16152 ± 1138 | 16006 ± 1609 | 0.94 |
| CA3 | 11837 ± 2436 | 10475 ± 2103 | 0.68 |

All data are presented as mean ± SEM, unpaired t-test.

**Supplementary Table 2**

Quantification of cholinergic ChAT positive neurons in the basal forebrain of sham (n = 5) and septic mice (n = 6)

| # of ChAT positive neurons | | | |
|---|---|---|---|
| Treatment | | | |
| Brain region | Sham (mean ± SEM) | Sepsis (mean ± SEM) | p-value |
| Basal forebrain | 24 ± 2.63 | 19.4 ± 3.04 | 0.28 |

All data are presented as mean ± SEM, unpaired t-test.

**Supplemental Figures:**

**Supplementary Figure 1.**

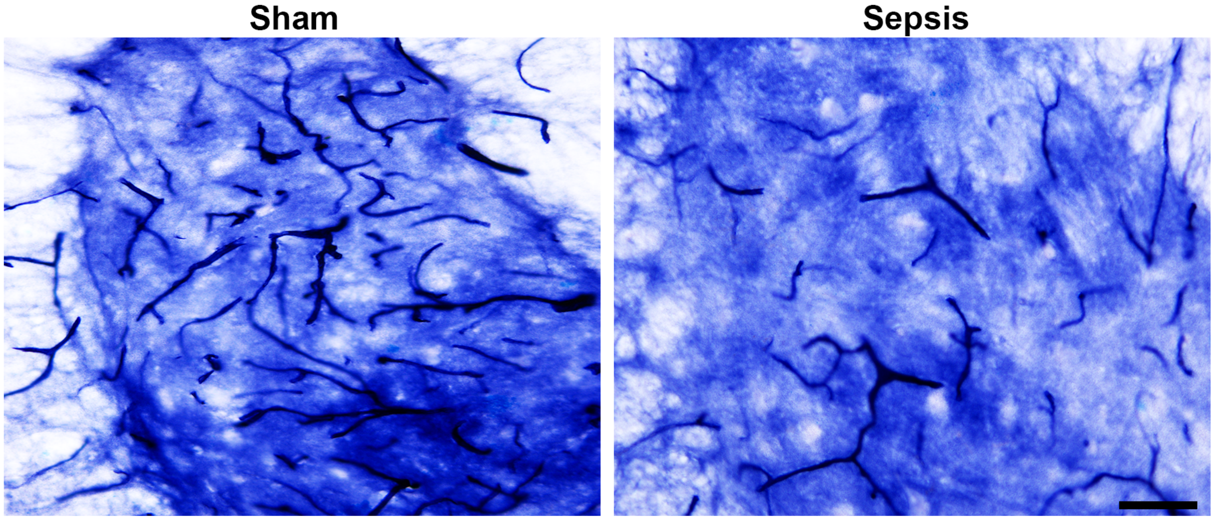

### Supplementary Figure 2.

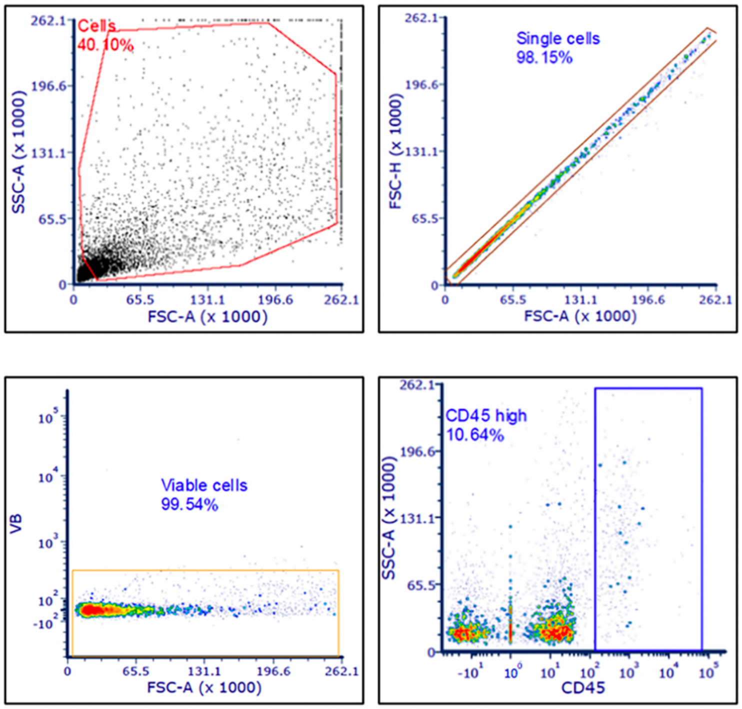

**Supplementary Figure 3.**

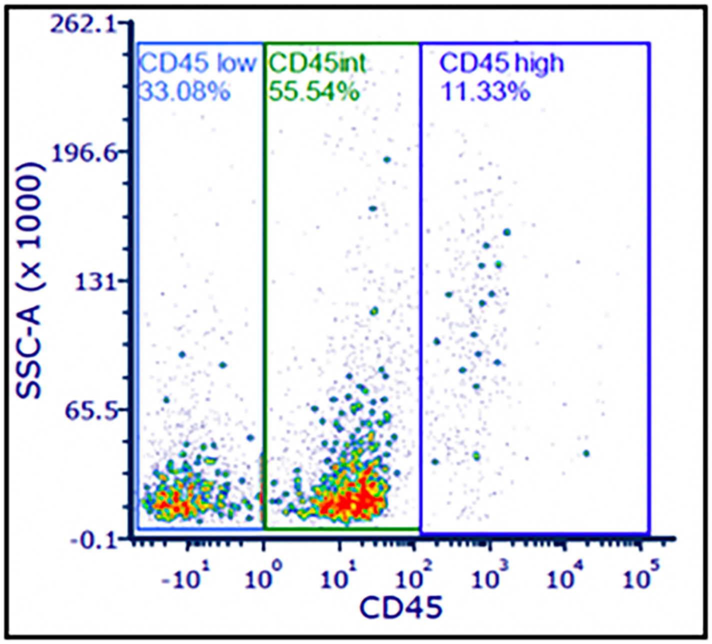

**Supplementary Figure 4.**

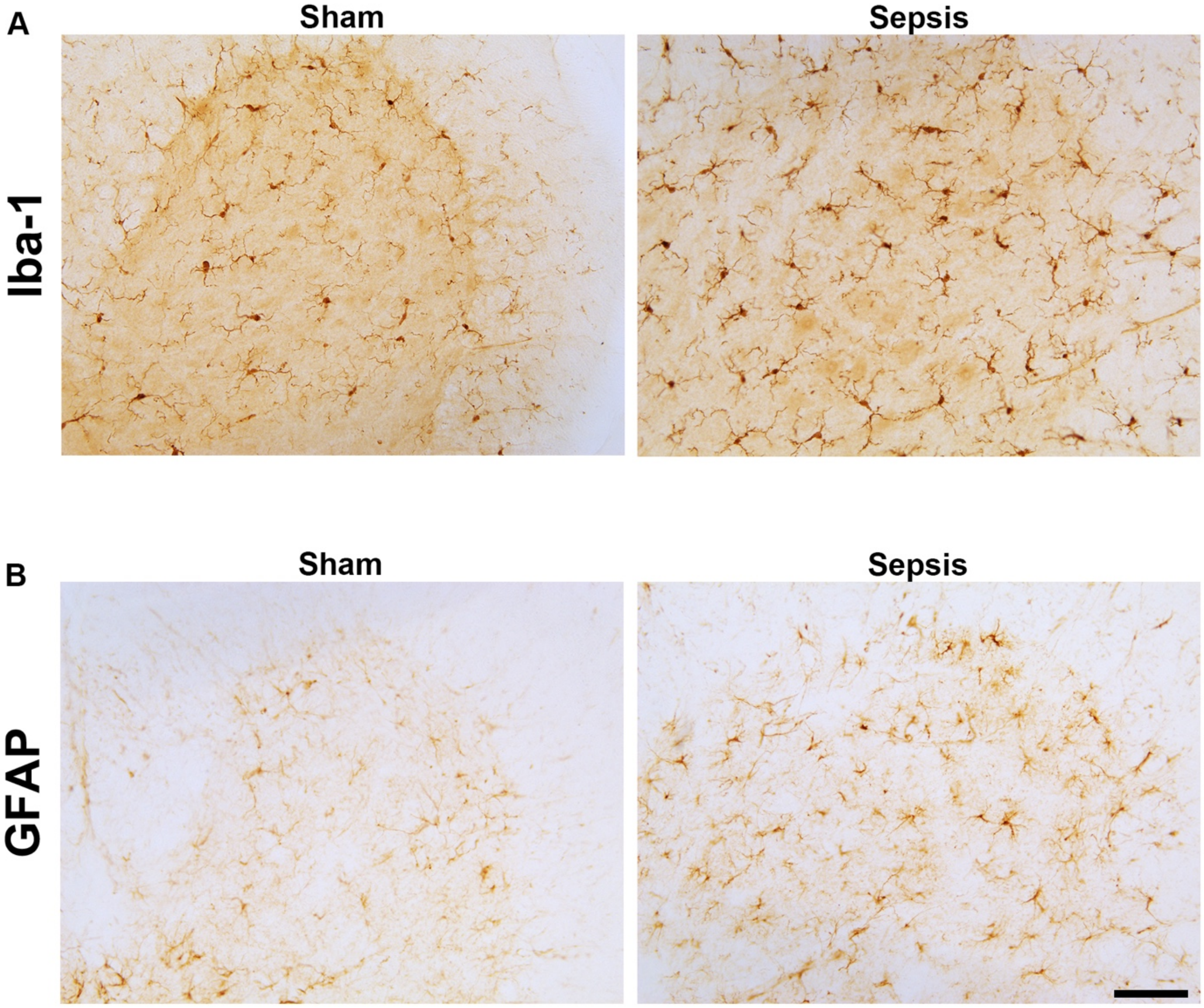

**Supplementary Figure 5.**

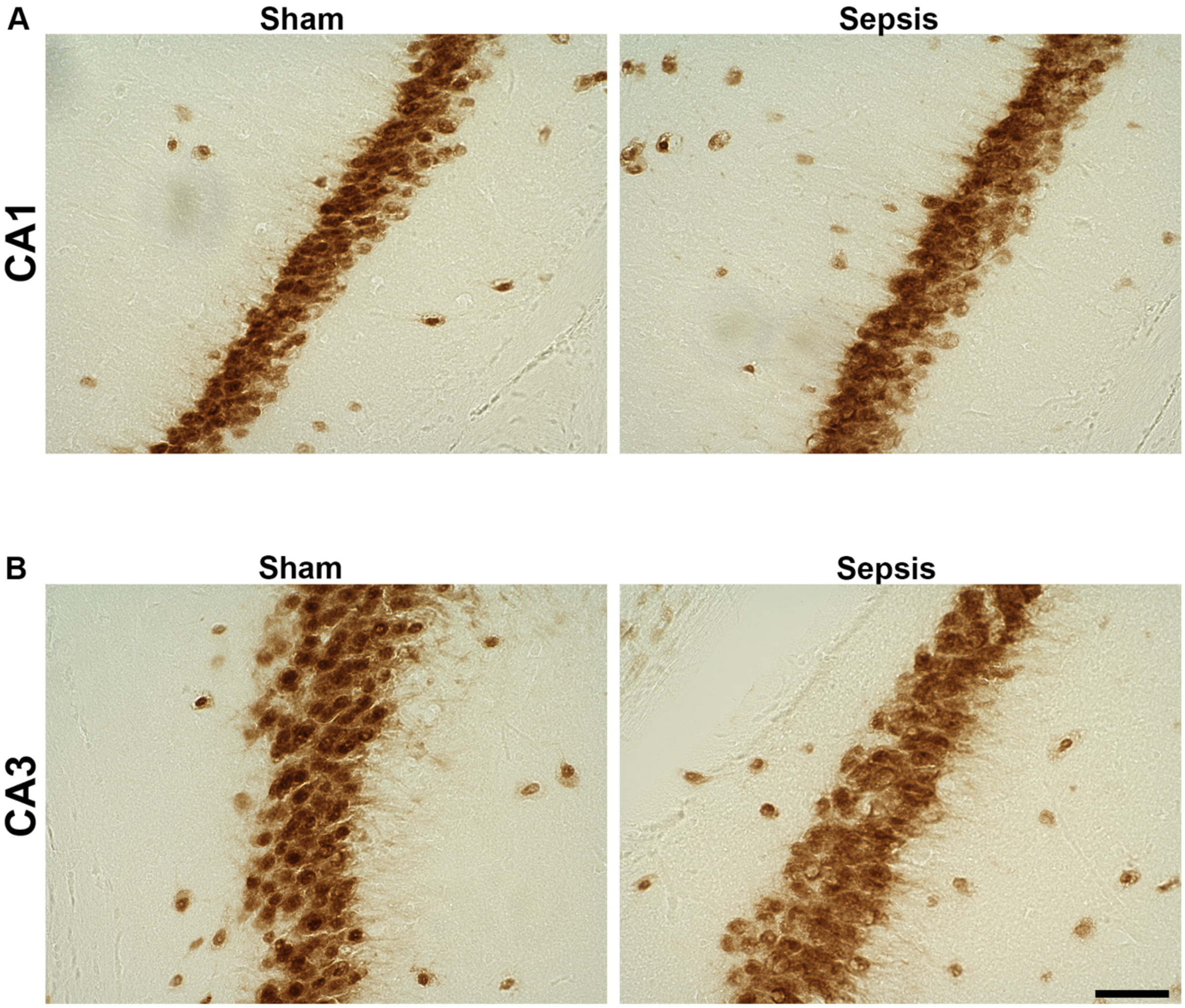

**Supplementary Figure Legends**

**Supplementary Figure 1.**

**Tissue non-specific alkaline phosphatase (TNAP) enzyme activity in the spinal cord.** Representative alkaline phosphatase (AP) histology shows that TNAP's enzyme activity is decreased in the spinal cord of septic mice compared to sham mice (20X magnification). 20X magnification scale bar = 80 μm.

**Supplementary Figure 2.**

**Gating strategy for brain myeloid and lymphoid cell populations.** All myeloid and lymphoid cell lineages used in our analyses are under the parent gate $CD45^{hi}$ prior to being organized into precise cell types using the following specific markers: CD11b, CD11c, Ly6G, Ly6C, CD3, CD4, CD8, CD25, and Foxp3. SSC = side scatter, FSC = forward scatter, A = area, H = height, and VB = viability. Analyzed population is indicated by the encircled gating.

**Supplementary Figure 3.**

**Gating strategy for brain microglia/monocyte populations.** The brain microglia/monocytes population used for analysis is encircled under the parent gate $CD45^{low}$. SSC = side scatter and A = area.

**Supplementary Figure 4.**

**Iba-1 positive microglia and GFAP positive astrocyte immunohistochemistry in the spinal cord of septic and sham mice.** (A) Iba-1 positive cells are increased in the spinal cords of septic mice compared to sham mice. (B) GFAP positive cells are increased in the spinal cords of septic mice compared to sham mice (20X magnification). 20X magnification scale bar = 80 μm.

**Supplementary Figure 5.**

**NeuN positive immunohistochemistry for neuronal loss in the CA1 and CA3 hippocampal regions.** Representative histology for NeuN immunolabelling shows no apparent differences in neuronal loss in the (A) CA1 or (C) CA3 between septic and sham mice (40X magnification). 40X magnification scale bar = 40 μm.
